# Stability of c-Myc protein differentiates Ras oncogene addiction and MAPK pathway dependency in Ras-mutant multiple myeloma

**DOI:** 10.64898/2026.08.06.743109

**Authors:** Yeon-Hwa Lee, Christophe Cataisson, Haibo Zhang, Snehal Gaikwad, Wendy D. du Bois, Aleksandra M. Michalowski, Howard H. Yang, Thomas J. Meyer, Ryan M. Young, Beverly A. Mock, Ji Luo

## Abstract

Multiple myeloma (MM) is a plasma cell malignancy that frequently harbors activating mutations in *NRAS* and *KRAS* oncogenes. Previous clinical trials targeting the Ras/MAPK oncogenic pathway with MEK inhibitors (MEKi) were met with limited efficacy, and newer generation of Ras inhibitors (RASi) have not been specifically evaluated in MM patients. To investigate the vulnerabilities of Ras-mutant MM to targeted therapies, we examined the sensitivity of a panel of human MM cell lines to the RASi RMC-6236 (daraxonrasib) and the MEKi trametinib. Although Ras-mutant MM cells are responsive to oncogenic Ras signaling and are sensitive to RAS inhibition, their sensitivity to MEK inhibition is heterogeneous. Mechanistic studies revealed that c-Myc protein is destabilized by MEK inhibition only in MEKi-sensitive MM cells but not in MEKi-resistant cells, and pharmacological and genetic stabilization of c-Myc is sufficient to confer MEKi resistance. In contrast, Ras inhibition reduced c-Myc protein across all MM cell lines tested, regardless of their dependency on the MAPK pathway, and c-Myc expression was insufficient to promote RASi resistance. Together, these findings demonstrate that c-Myc protein stability differentiates the response of Ras-mutant MM cells to Ras and MEK inhibition, and suggest that direct targeting of the Ras oncoprotein, rather than its downstream MAPK pathway, may present a more effective strategy.

## INTRODUCTION

Multiple myeloma (MM) is a plasma cell malignancy and the second most common hematologic cancer in the United States (1). About 40% of MM cases harbor chromosomal translocations between the immunoglobulin heavy chain (IgH) locus and other genes including *CCND1* and *MYC* that result in cyclin D1 and c-Myc over-expression, respectively, to promote cell proliferation and survival (2–4). MM is also characterized by frequent mutations in the Ras/MAP kinase (MAPK) pathway including activating mutations in *KRAS* (∼23%), *NRAS* (∼20%), and less frequently, *BRAF* (∼6%) (5–7). These mutations are less frequent in the precursor conditions monoclonal gammopathy of undetermined significance (ranging from 0.5-2.7%) and smoldering multiple myeloma (1.4%-14.8%) (8). Ras/RAF mutations are detectable in approximately half of newly diagnosed MM cases, and more common in relapsed and drug-resistant MM (5–7). Thus, Ras oncogene is a key driver of MM progression and a promising therapeutic target. The MAPK pathway, composed of the RAF-MEK-ERK kinase cascade, is a central regulator of cell proliferation and survival downstream of Ras. Activation of this pathway by the Ras oncoprotein promotes cell cycle progression, in part, by increasing the transcription of D-type cyclins and c-Myc (9, 10). ERK can directly phosphorylate c-Myc and enhance c-Myc stability by preventing its proteasomal degradation (11, 12). Oncogenic transformation driven by the Ras/MAPK pathway is interconnected with c-Myc function (13). Several studies have shown that high c-Myc expression is both critical for the viability of MM cells and can promote their drug resistance (14–16). The co-existence of *MYC* translocation/rearrangement and *Ras* mutation in MM strongly suggest that c-Myc and Ras/MAPK could cooperate to promote MM progression.

Advances in MM treatment, including proteasome inhibitors, immunomodulatory drugs, and monoclonal antibodies, have significantly improved the survival of MM patients (17). However, the 10-year survival rate for MM patients remains only about 33% (18). Thus, new therapies are needed to improve durable disease control and extend the duration of clinical benefit. Pre-clinical studies showed that Ras mutant MM cells have elevated MAPK pathway activity (19). Clinical studies targeting the MAPK pathway in MM, however, had mixed outcomes. In a phase II trial of the MEKi selumetinib in treatment-refractory MM, partial response was observed in 2 of 36 patients, both of whom harbored Ras mutations (20). Similarly, a phase I trial of the RAF-MEK inhibitor CH5126766 reported one partial response among 6 MM patients with Ras/RAF mutations (21). These findings are consistent with the observation that Ras-mutant MM cell lines display variable MAPK pathway activation (22) and variable sensitivity to the MEK inhibitor trametinib (23), and they raise question on whether the MAPK pathway is a good target in Ras mutant MM. Recent development of inhibitors directly targeting the Ras oncoprotein have transformed treatment of lung and pancreatic cancer with KRAS mutation (24). Specific targeting of Kras in myeloma with AZD4785, a potent antisense oligonucleotide and tool compound that downregulates all Kras isoforms, inhibited MM cell growth, demonstrating its potential as a therapeutic target (24). A particularly promising new class of RASi are the multi-Ras(ON) inhibitors, exemplified by RMC-6236 (daraxonrasib), that can target all three Ras paralogs bearing different point mutations (25). The pre-clinical activity of RMC-6236 (25) and its efficacy in pancreatic cancer in a recent phase III trial (26) suggest that it could be a promising therapy for MM with Ras mutation.

In this study, we investigated the mechanisms underlying the sensitivity of Ras-mutant MM cells to the MEKi trametinib and the RASi RMC-6236 to evaluate their therapeutic potential. We found that c-Myc protein stability is a key determinant of cellular sensitivity to MEK inhibition. In contrast, despite their heterogeneous responses to MEK inhibition, Ras-mutant MM cells exhibited a more uniform sensitivity to direct Ras inhibition, suggesting that RAS inhibitors could provide greater therapeutic benefit.

## RESULTS

### Ras mutant MM cells exhibit Ras oncogene addiction but heterogeneous MAPK pathway dependency

To investigate the sensitivity of Ras mutant MM cells towards inhibitors targeting the Ras/MAPK pathway, we assembled a panel of Ras mutant cell lines including 4 NRAS mutant cells lines INA6 (heterozygous NRAS^WT/G12D^), NCI-H929 (heterozygous NRAS^WT/G13D^, hereafter referred to as H929), JJN3 (heterozygous NRAS^WT/Q61K^) and L363 (hemizygous NRAS^Q61H^), and 2 KRAS mutant cell lines RPMI8226 (heterozygous KRAS^WT/G12A^) and KMS28BM (heterozygous KRAS^WT/G12A^). To evaluate the functional addiction of Ras-mutant MM cells on Ras signaling for proliferation, we transduced MM cells with lentiviral shRNAs targeting NRAS and KRAS. Knocking down the Ras oncogene led to a reduction in ERK1/2 kinase (hereafter referred to as ERK) phosphorylation, indicating downregulation of MAPK pathway activity (**Fig. 1A**). Concomitantly, cell viability was significantly decreased (**Fig. 1B**), confirming that these Ras mutant MM cell lines are dependent on the Ras oncogene for MAPK pathway signaling and for proliferation. Data mining using the DepMap database (https://depmap.org/portal/) (27) independently confirmed that these cell lines are sensitive to CRISPR-mediated knockout of their respective Ras oncogene (**Fig. S1A**). The INA6 cell line, which harbors *CCND1* translocation, has the highest expression of cyclin D1 and is sensitive to *CCND1* knockout. Other cell lines show higher expression of cyclin D2 and are sensitive to *CCND2*, but not *CCND1*, knockout, consistent with cyclin D2 being a key D-type cyclin in the B-cell lineage (28, 29). The expression of c-Myc is high in all cell lines regardless of their MYC gene translocation status, and all cell lines are sensitive to MYC knockout (**Fig. S1A**).

**Figure 1.**
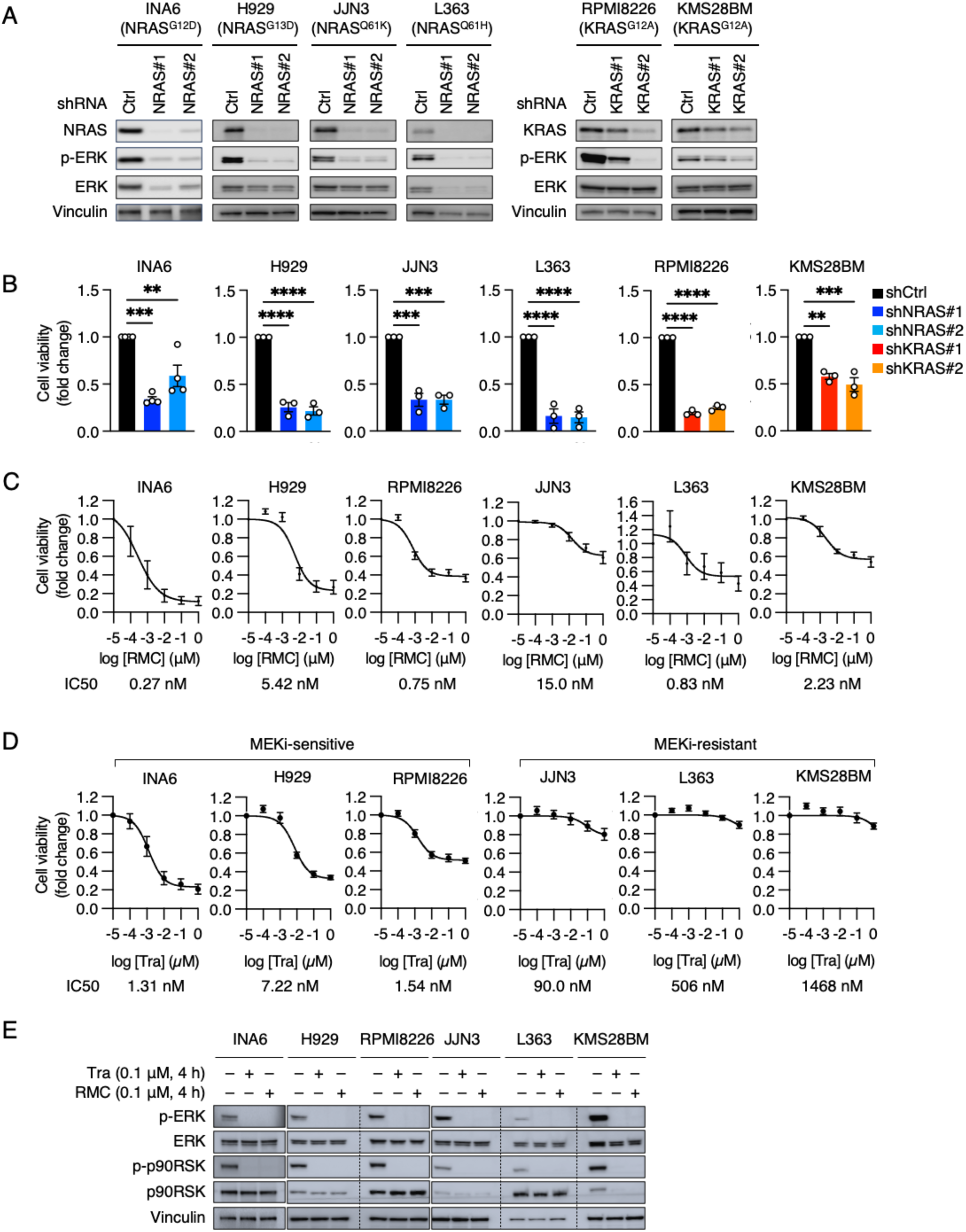
Ras-mutant MM cells are sensitive to Ras inhibition but show variable sensitivity to MEK inhibition. A. Inhibition of MAPK pathway by Ras knockdown. Human MM cell lines were transduced with either control (Ctrl) lentiviral shRNA or shRNAs targeting *NRAS* and *KRAS*. Ras protein knockdown and phospho-ERK level (p-ERK) was evaluated by immunoblot 5 days post-transduction. B. Inhibition of cell viability by Ras knockdown. Cells were transduced with shRNAs and cell viability was measured 3 days after plating using CellTiter-Glo assay (unless otherwise specified, for all figure panels statistical significance is annotated as * *p* < 0.05, ** *p* < 0.01, *** *p* < 0.001, and **** *p* < 0.0001, and error bars represent mean ± SEM of ≥ 3 independent experiments). C. Effect of RASi on cell viability. Cells were treated with the Multi-Ras(ON) inhibitor RMC-6236 (RMC) for 4 days and cell viability was measured using CellTiter-Glo assay. IC_50_ value represents drug concentration at 50% maximal effect. D. Effect of MEKi on cell viability. Cells were treated with the MEK inhibitor trametinib (Tra) for 4 days and cell viability was measured using CellTiter-Glo assay. E. MAPK pathway inhibition by RASi and MEKi. Cells were treated with trametinib and RMC-6236 for 4 hours and cell lysates were immunoblotted for p-ERK and phospho-p90RSK (p-p90RSK) to assess MAPK pathway inhibition.

To validate Ras dependency pharmacologically, we examined the sensitivity of these cell line to the multi-Ras(ON) inhibitor RMC-6236 (daraxonrasib), which can block both mutant NRAS and KRAS (25). All cell lines showed low nM IC_50_ values towards RMC-6236, although the maximum degree of growth inhibition was variable (**Fig. 1C, Fig S1B**), indicating that they retain substantial Ras dependence. Unexpectedly, when we examined the sensitivity of these cell lines towards trametinib, a selective MEK1/2 kinase inhibitor, we found that they exhibited heterogeneous sensitivity to MEK inhibition. Three cell lines, INA6, H929, and RPMI8226 are MEKi-sensitive with low-nM IC_50_ values. In contrast, JJN3, L363, and KMS28BM cells are MEKi-resistant with minimal growth inhibition at dose ranges limiting off-target effects (**Fig. 1D, Fig S1B**). Both RMC-6236 and trametinib were effective at fully blocking ERK phosphorylation at sub-μM concentrations (**Fig. 1E**). Thus, the difference in MEKi sensitivity is not likely attributable to difference in MAPK pathway inhibition by trametinib. Neither is MEKi sensitivity correlated with the proliferation rate of these cell lines (**Fig. S1C and S1D**). Data mining from a previous drug sensitivity study on a large panel of MM cell lines (30) showed that NRAS and KRAS mutant MM cells trended to be more sensitive to trametinib than Ras WT cells, although Ras mutant cells exhibit a wide range of trametinib IC_50_ values (**Fig. S1E**). Correlating trametinib IC_50_ data with Ras oncogene CRISPR dependency scores (27) revealed a weak correlation that did not reach significance (**Fig. S1F**). These independent datasets therefore support the notion that Ras oncogene dependency and MEKi sensitivity may have distinct attributes. Importantly, both MEKi-resistant and - sensitive cells are comparably sensitive to the proteasome inhibitor bortezomib and the chemotherapeutic agent doxorubicin (**Fig. S1G and S1H**). Thus, MEKi-resistance is not likely due to broad-spectrum drug resistance mechanism. Together, these results indicate that Ras-mutant MM cells exhibit addiction to the Ras oncogene but heterogeneous dependency on the MAPK pathway.

### MEKi and RASi differentially regulate D-type cyclins and c-Myc levels in MM cells

To understand the functional differences between MEKi-sensitive and-resistant cells, we first examined the effect of MEK inhibition on cell proliferation. In MEKi-sensitive cells, trametinib led to G1 arrest, while having only minimal effect on the proliferation of MEKi resistant cells (**Fig. S2A**). Trametinib did not induce significant cell death, as evaluated by May-Grünwald Giemsa staining (**Fig. S2B**) or by immunoblotting of cleaved caspase-3 and cleaved PARP protein (**Fig. S2C**). This lack of apoptosis is not due to a defect in apoptotic response, as doxorubicin readily induced apoptosis in all cell lines (**Fig. S2C**). Thus, MEK inhibition primarily induces growth arrest rather than cell death in MEKi-sensitive cell lines.

To identify molecular changes that are associated with MEKi sensitivity, we examined the effect of trametinib on a selected group of protein and phosphoproteins, including those in the MAPK and AKT/mTOR signaling pathway and in cell cycle regulation, that are known to mediate the proliferation effects of the Ras. Immunoblots showed that these analytes fall into three categories. The first category is composed of analytes that are down-regulated by trametinib in both MEKi-sensitive and MEKi-resistant cell lines. These include phospho-ERK (p-ERK), phospho-p90RSK (p-RSK, which is a direct ERK phosphorylation target), and DUSP4 and DUSP6, two phosphatases induced by ERK signaling (**Fig. 2A, Fig. S3A and S3B**). The second category is analytes that are down-regulated by trametinib only in MEKi-sensitive but not in MEKi-resistant cell lines. These include c-Myc, phospho-S6K kinase (p-S6K) and phospho-ribosomal S6 protein (p-S6), and cyclins D1 and D2 (**Fig. 2A, 2B and Fig. S3A, S3C**). We carried out Spearman’s correlation analysis of the area under the curve (AUC) values of trametinib and AUC values of these analytes (**Figure S3E**). We saw a strong, positive correlation between trametinib sensitivity and trametinib-induced changes in c-Myc, cyclin D2, and p-S6 levels (**Fig. 2C, Fig. S3F**). The third category is analytes that are not downregulated by trametinib in any of the cell lines. These include phospho-AKT (p-AKT) and cyclin D3 (**Fig. 2A, Fig. S3A and S3D**).

**Figure 2.**
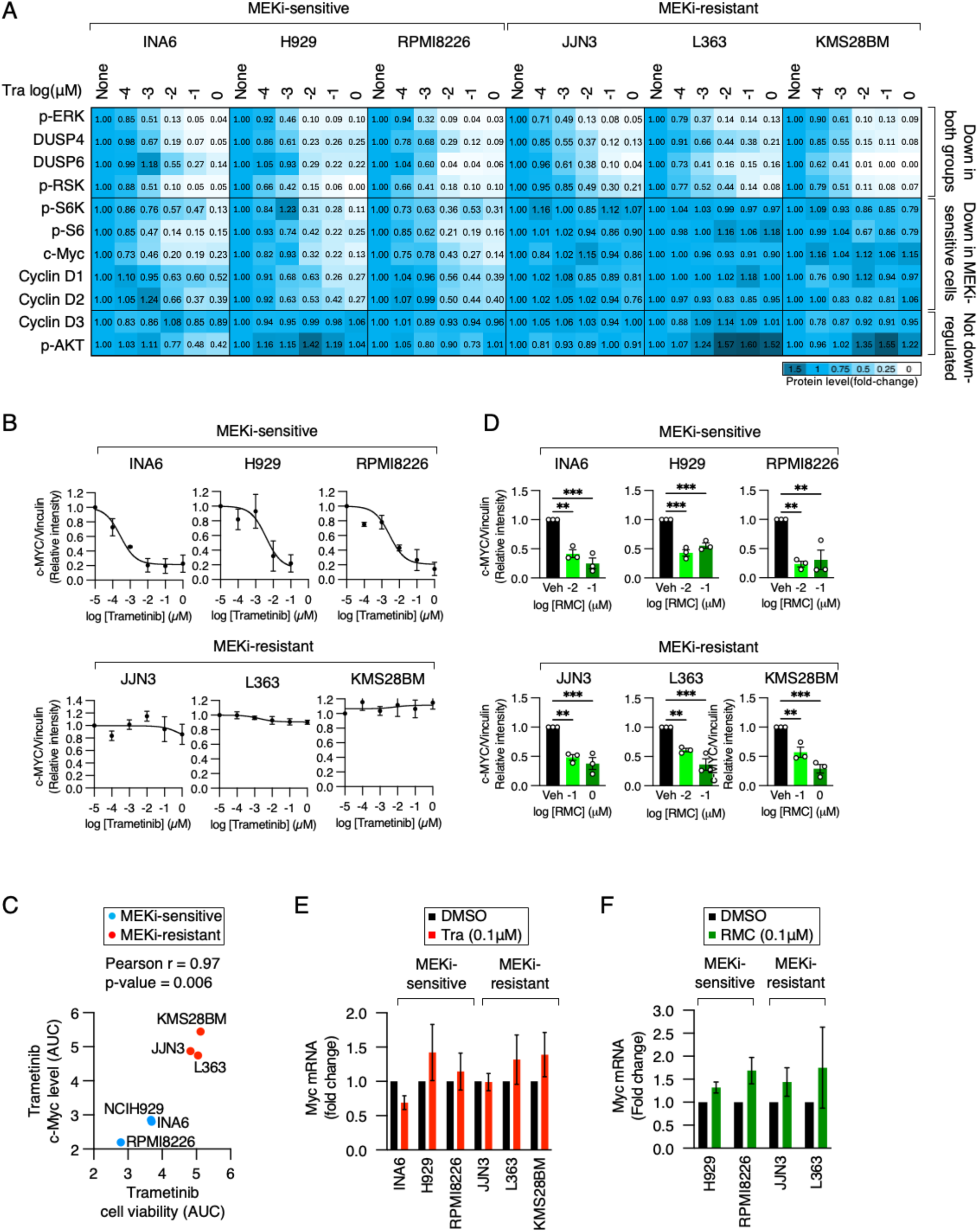
MEK and Ras inhibition differentially regulate c-Myc protein level. A. Heatmap summarizing changes in protein analyte levels in response to MEKi. Cells were treated with trametinib for 4 days and cell lysates were immunoblotted for the indicated proteins and phospho-proteins. Protein signal was quantified by densitometry. B. Effect of MEKi on c-Myc protein level. Cells were treated with trametinib for 4 days and dose-response curve of c-Myc protein level was generated by immunoblotting. C. Correlation between cell viability and c-Myc protein changes induced by MEKi. AUC values of cell viability and c-Myc protein level in response to trametinib are evaluated by Pearson correlation. D. Effect of RASi on c-Myc protein level. Cells were treated with RMC-6236 for 4 days and c-Myc protein level was quantified by immunoblotting. E. Effect of MEKi on c-Myc transcript level. Cells were treated with trametinib for 4 days and c-Myc mRNA level was quantified by RT-qPCR. F. Effect of RASi on c-Myc transcript level. Cells were treated with RMC-6236 for 4 days and c-Myc mRNA level was quantified by RT-qPCR.

Next, we examined how RASi affects the level of c-Myc, cyclin D2 and p-S6 in these cells. Interestingly, RMC-6236 treatment reduced the level of these proteins in all cell lines regardless of their MEKi sensitivity status (**Fig. 2D, Fig. S4**). Taken together, these results indicate that MEKi resistance is not due to the inability of trametinib to block MAPK signaling, and the difference in MEKi and RASi sensitivity among MM cells is likely associated with the differential regulation of c-Myc, cyclin D2, and p-S6 levels by these inhibitors.

### c-Myc protein stability confers resistance to MEKi but not to RASi

Given that MEKi sensitivity is associated with down-regulation of c-Myc, and c-Myc expression has been shown to drive resistance to MAPK pathway inhibition in solid tumor cell lines (31), we tested the hypothesis that sustained c-Myc protein expression can drive MEKi resistance in MM cells. Measurement of c-Myc mRNA level showed no down-regulation of c-Myc expression in response to trametinib in all cell lines (**Fig. 2E**). Similarly, c-Myc mRNA levels were not down-regulated by RMC-6236 treatment in two MEKi-sensitive and two MEKi-resistant cell lines (**Fig. 2F**). This suggests that the loss of c-Myc protein in response to these inhibitors is not due to its transcriptional downregulation but rather its protein degradation. As c-Myc is a short-lived protein (32), we measured the half-life of c-Myc protein in MM cells using cycloheximide chase assays. The degradation of c-Myc protein was significantly accelerated by trametinib treatment in MEKi-sensitive cells but not in MEKi-resistant cells (**Fig. 3A and Fig. S5A**). In MEKi-sensitive cells, the baseline half-life of c-Myc was somewhat longer than that in MEK-resistant cells, and c-Myc half-life was reduced by approximately 50% upon trametinib treatment. In contrast, the half-life of c-Myc was unchanged by trametinib treatment in MEKi-resistant cells (**Fig. 3B)**. Pearson correlation analysis confirmed a strong correlation between the reduction in c-Myc half-life and trametinib AUC (**Fig. 3C**).

**Figure 3.**
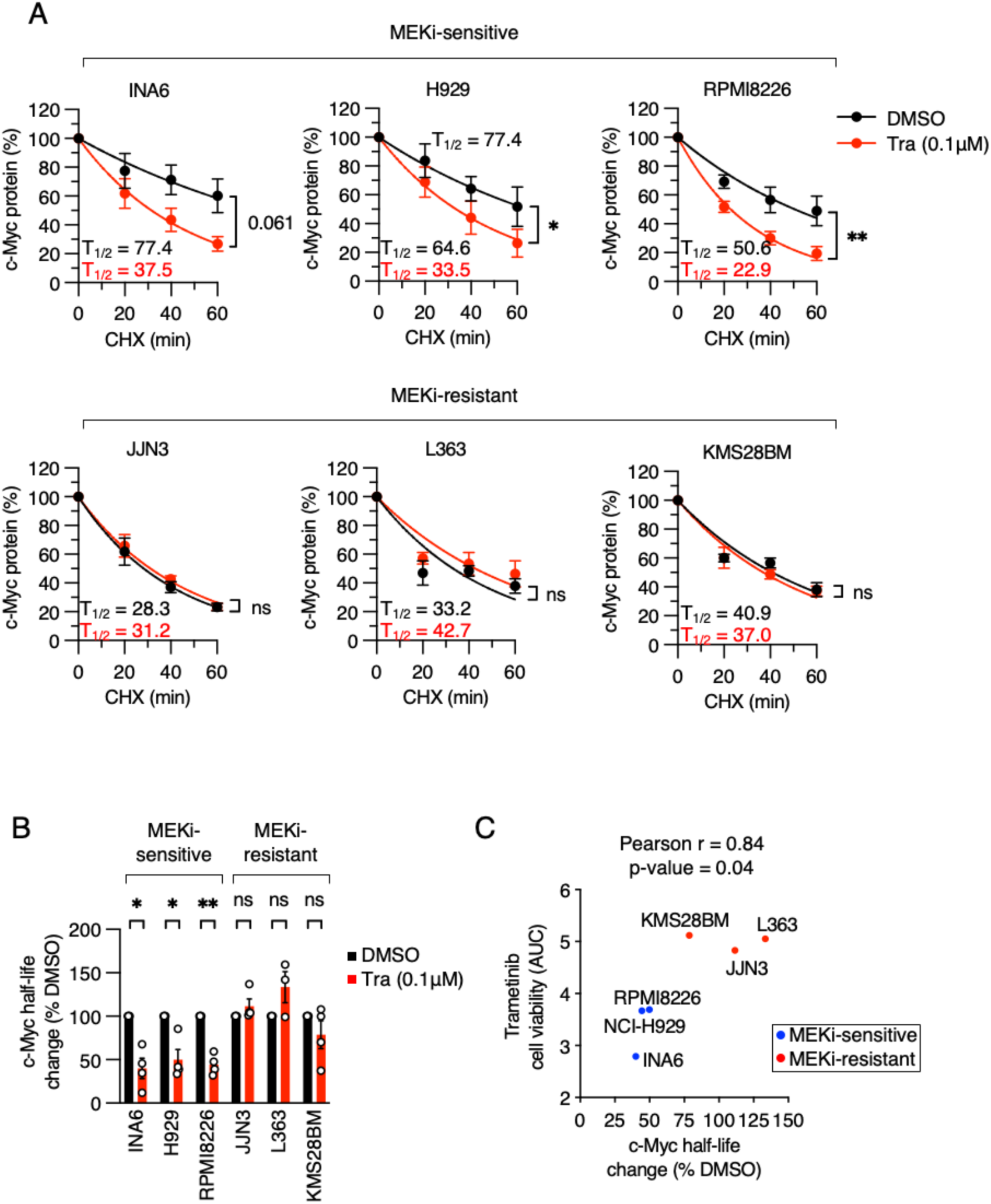
MEK inhibition destabilizes c-Myc protein in MEKi-sensitive MM cells. A. Effect of MEKi on c-Myc protein stability. Cells were treated with trametinib for 4 hours followed by cycloheximide (CHX) co-treatment for the indicated time to block protein translation. The level of c-Myc protein was measured by immunoblotting and quantified by densitometry. Half-life of c-Myc was estimated by fitting to a one-phase decay curve. B. Changes in c-Myc protein half-life in response trametinib. Half-life of c-Myc in trametinib-treated cells were normalized to that in vehicle (DMSO) treated cells. C. Correlation analysis between trametinib cell viability AUC and chang in c-Myc protein half-life.

The stability of c-Myc protein is regulated by a series of phosphorylation/dephosphorylation events. Upon growth stimulation, ERK and CDKs have been shown to phosphorylate S62 on c-Myc to both stabilize it and prime it for phosphorylation by GSK3β at T58 when growth signals are reduced (11). This leads to subsequent dephosphorylation at S62 by PP2A creating a phospho-degron signal that targets c-Myc for ubiquitination by FBXW7 and proteasomal degradation (33). We therefore examined whether c-Myc degradation could be causal to MEKi sensitivity. In MEKi-sensitive cells, trametinib treatment led to the rapid shutdown of ERK activity as indicated by the loss of both phosphorylated ERK and phosphorylated p90RSK downstream of ERK within one hour. The level of S62 phosphorylated c-Myc, on the other hand, declined very slowly in 24 hours and was largely correlated with total c-Myc protein level (**Fig. S5B and S5C**). This indicates that either the dephosphorylation of S62 is very slow following ERK inactivation, or other proline-directed kinases may also phosphorylate S62 on c-Myc independent of ERK in MM cells. To stabilize c-Myc, we treated cells with a GSK3β-selective inhibitor CHIR99021. This inhibitor at 1 μM led to a decrease in T58 phosphorylated c-Myc and a corresponding increase in total c-Myc level (**Fig. S5D**). In MEKi-sensitive cells, co-treatment of CHIR99021 stabilized c-Myc level in the presence of trametinib, thus counteracting its destabilizing effect (**Fig. 4A and Fig. S5E**). Functionally, CHIR99021 decreased the sensitivity of these cells to trametinib (**Fig. 4B**). Next, we constitutively expressed either wild type c-Myc or the c-Myc T58A mutant which cannot be phosphorylated by GSK3β (11) in the MEKi-sensitive RPMI8226 cell line. This approach elevated total c-Myc protein level in the presence of trametinib (**Fig. 4C**) and rendered these cells more resistant to trametinib (**Fig. 4D**). Thus, elevating c-Myc protein level, through either pharmacological or genetic means, can confer resistance to MEKi. To further support this hypothesis, we inhibited c-Myc by expressing omoMyc under a tetracycline-inducible promoter in the MEKi-resistant JJN3 cell line (**Fig. 4E**). omoMyc is a mini-peptide derived from c-Myc that blocks Myc-MAX dimerization and therefore c-Myc function (34). Despite its relatively low expression upon doxycycline induction, JJN3 cells expressing omoMyc trended towards increased trametinib sensitivity, although the p-value did not reach statistical significance (**Fig. 4F**). Taken together, these results support the conclusion that c-Myc stability can be a key regulator of MEKi sensitivity in Ras mutant MM cells.

**Figure 4.**
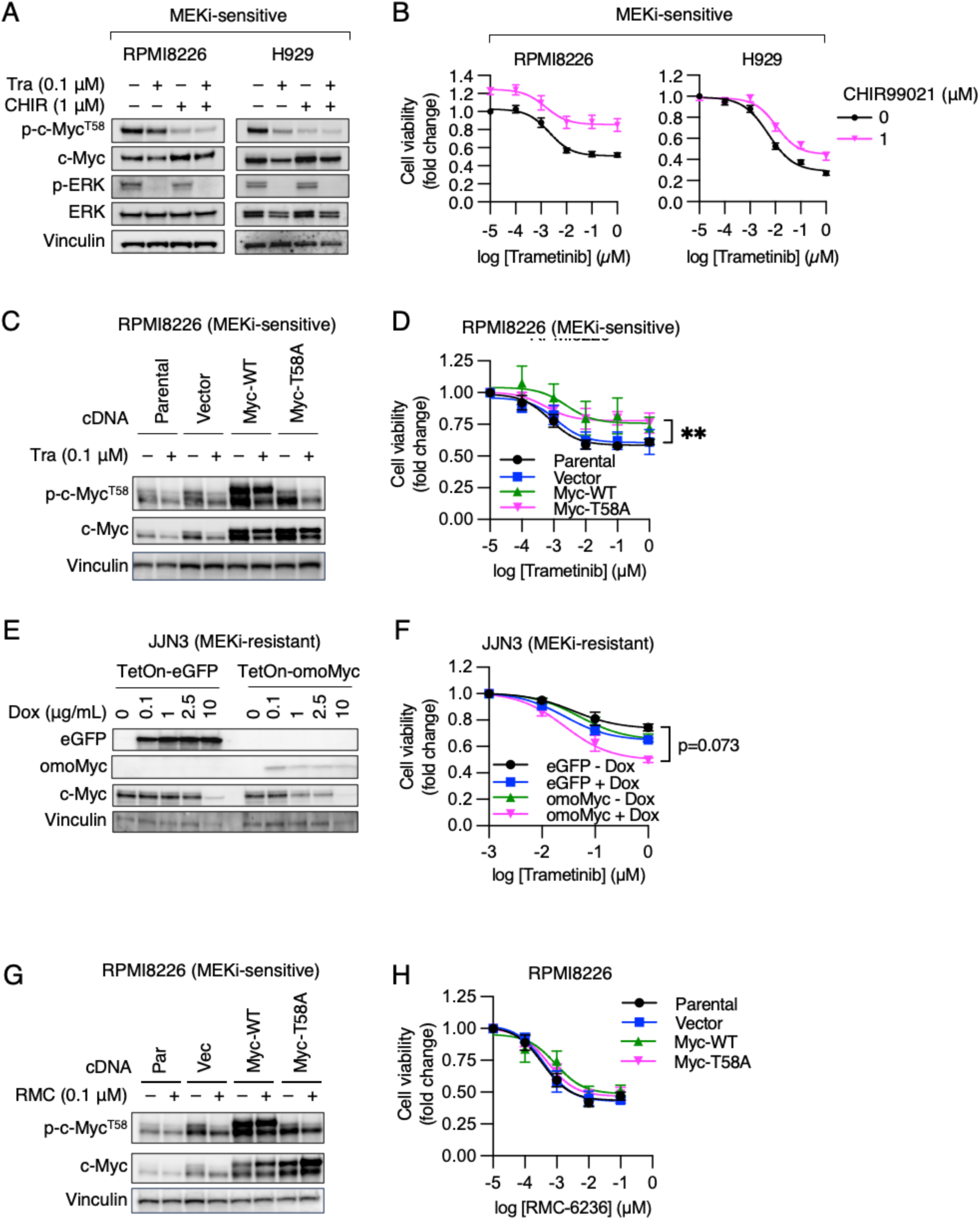
c-Myc stabilization confers MEKi resistance but not RASi resistance. A. Effect of GSK3β inhibition on c-Myc stability. MEKi-sensitive RPMI8226 and H929 cells were treated with trametinib and the GSK3β inhibitor CHIR99021 either alone or in combination for 24 hours. Changes in phospho-c-Myc^T58^ and total c-Myc protein level were analyzed by immunoblotting. B. Effect of GSK3β inhibition on MEKi sensitivity. Trametinib dose-response of MEKi-sensitive RPMI8226 and H929 cells were measured with or without combination treatment with CHIR99021. Cells were treated with inhibitors for 4 days and cell viability was assessed using CellTiter-Glo assay. C. Overexpression of exogenous c-Myc. RPMI8226 cells were transduced with lentiviral vector stably expressing FLAG-tagged wild type c-Myc (Myc-WT) or T58A mutant c-Myc (Myc-T58A). Total c-Myc and p-c-Myc T58 with or without trametinib treatment were analyzed by immunoblotting (Par, parental cells; Vec, vector only control). D. Effect of c-Myc overexpression on MEKi sensitivity. MEKi-sensitive RPMI8226 cells expressing Myc-WT or Myc-T58A were treated with trametinib for 4 days and cell viability was assessed using CellTiter-Glo assay. E. Expression of dominant-negative omoMyc. JJN3 cells were stably transduced with lentiviral vectors expressing tetracycline-inducible (tet-on) HA-tagged cDNAs of eGFP or omoMyc. Protein expression was tested at indicated doses of doxycycline (Dox). F. Effect of omoMyc expression on MEKi sensitivity. MEKi-resistant JJN3 cells with (+Dox) or without (-Dox) eGFP or omoMyc expression were treated with trametinib for 4 days and cell viability was assessed using CellTiter-Glo assay. G. Overexpression of exogenous c-Myc. RPMI8226 cells were transduced with lentiviral vector stably expressing FLAG-tagged wild type c-Myc (Myc-WT) or T58A mutant c-Myc (Myc-T58A). Total c-Myc and p-c-Myc T58 with or without RMC-6236 treatment were analyzed by immunoblotting (Par, parental cells; Vec, vector only control). H. Effect c-Myc overexpression on RASi sensitivity. RPMI8226 cells expressing Myc-WT or Myc-T58A were treated with RMC-6236 for 4 days and cell viability was assessed using CellTiter-Glo assay.

As the RASi RMC-6236 was able to down-regulate c-Myc protein in MEKi resistant cells, we asked the question whether c-Myc overexpression could also confer resistance to RMC-6236. Neither wild type nor T58A c-Myc was able to promote resistance to RMC-6236 in RPMI8226 cells (**Fig. 4H**), despite robust overexpression of c-Myc, as confirmed by immunoblotting **(Fig. 4G)**. Thus, whereas c-Myc can overcome the dependence of MM cells on the MAPK pathway, it was insufficient to overcome their dependence on the Ras oncoprotein.

### Cyclin D2, AKT, mTOR/S6K signaling, and noncanonical MAPK do not explain MEKi resistance

In addition to c-Myc, we examined whether Cyclin D2, AKT, mTOR/S6K signaling, or noncanonical MAPK can contribute to MEKi resistance in MM cells. In MM cells, a key Ras effector is mTOR (35). Since S6 phosphorylation correlated with MEKi sensitivity, a plausible hypothesis is that Ras-sustained signaling through the mTOR/S6K axis can drive MEKi resistance. We tested whether inhibiting S6K kinase with the inhibitor LY2584702 (36) can sensitize MEKi-resistant cells towards trametinib. LY2584702 effectively blocked S6 phosphorylation and showed robust single-agent activity in MEKi-resistant cells. However, it did not sensitize them to trametinib in a combination setting (**Fig. S6A and S6B**). Similarly, the mTORC1 inhibitor rapamycin displayed variable single-agent activity in MEKi-resistant cells but did not sensitize them to trametinib (**Fig. S6C**). As the PI3K/AKT signaling pathway have been shown to reduce the dependence of Ras mutant cells on MAPK pathway in solid tumor cells (37, 38), we blocked AKT signaling using the selective inhibitor MK-2206. While MK-2206 displayed robust single-agent activity in MEKi resistant cells, it did not sensitize them to trametinib (**Fig. S6D**). Thus, both mTOR/S6K signaling and PI3K/AKT signaling are unlikely to contribute to MEKi resistance in this context.

Cyclin D2 is an important regulator of cell cycle entry in hematopoietic cells (39, 40). Like cyclin D2 protein, cyclin D2 mRNA was downregulated by trametinib in MEKi-sensitive cells but not in MEKi-resistant cells (**Fig. S7A**). We therefore tested whether cyclin D2 can confer MEKi resistance by overexpressing cyclin D2 from a tetracycline-inducible promoter in MEKi-sensitive H929 cells (**Fig. S7B**). The approach compensated the loss of endogenous cyclin D2 upon trametinib treatment **(Fig. S7C**), but it did not confer resistance to trametinib (**Fig S7D**). As an alternative approach, we tested whether blocking the G1 cyclin-dependent kinases CDK4/6 using palbociclib (41) can confer MEKi sensitivity. Palbociclib showed robust single-agent activity in MEKi-resistant cells but did not sensitize them towards trametinib (**Fig. S7E**). Thus, sustained cyclin D2 activity is insufficient to promote MEKi resistance.

Previous report has shown that alternative, noncanonical MAPK signaling through MEK5 and ERK5 contributes to ERK inhibitor resistance in KRAS mutant pancreatic cancer cells (31). We therefore tested whether the MEK5-ERK5 axis could promote trametinib resistance. Treatment of MEKi-resistant cells with the selective MEK5 inhibitor BIX02189 (42) and the selective ERK5 inhibitor XMD8-92 (42) showed various degree of single-agent activity. However, neither inhibitor, when combined with trametinib, sensitized these cells towards trametinib (**Fig. S8A and S8B**). We next examined whether other ERK family kinases including p38 and JNK could promote MEKi resistance. Treatment of MEKi-resistant MM cells with the selective p38α/β kinase inhibitor LY2228820 (43) and the selective JNK1/2/3 kinase inhibitor JNK inhibitor VIII (44) did not sensitize MEKi-resistant cells towards trametinib (**Fig. S8C and S8D**). Thus, non-canonical ERK kinase signaling is unlikely to contribute to MEKi resistance in these MM cells.

### Ras and MEK inhibitors elicit overlapping gene expression changes in MM cells

To understand the effect of Ras and MEK inhibition on gene-expression changes in Ras mutant MM cells, we performed RNA-seq analysis in two MEKi-sensitive cell lines, NCI-H929 and RPMI8226, and two MEKi-resistant cell lines, JJN3 and L363, after treatment with RMC-6236 and trametinib for 4 days to understand the steady-state gene expression changes elicited by stable MEK and Ras inhibition. Principle component analysis (PCA) showed that a major driver of gene expression pattern is cell-line features, whereas drug treatment contributed to a minor component in gene expression differences (**Fig. 5A**). Gene set enrichment analysis (GSEA) revealed a more pronounced down-regulation of hallmark pathways associated with cell proliferation, including E2F target, G2M check point, and mitotic spindle, in MEKi sensitive cells (**Fig. 5B**). Both drugs effectively blocked MAPK signaling (**Fig. 1E**). We saw strong down-regulation of a set of previously reported core ERK-dependent genes (45), including *DUSP6*, *ETV1/4/5*, *SPRY4* and *FOSL1,* in all four cell lines in response to both inhibitors (**Fig. 5C**). Genes commonly up-regulated by these two inhibitors are *RND3*, which encodes the small GTPase and negative cell cycle regulator RhoE (46), and *CELF6*, which encode an RNA binding protein with tumor suppressor activity (47). In MEKi-sensitive cells, a significant number of genes are up-or down-regulated by RAS and MEK inhibition. Among the down-regulated genes are those involved in cell cycle progression, including *CDC25A/C*, *AURKA*, *CENPA/E* and *MCM5/7* (**Fig. 5D**). In contrast, a much smaller set of genes were up-or down-regulated by these inhibitors in MEKi-resistant cells (**Fig. 5E**). In addition to these shared gene expression changes, we saw robust, cell-line specific gene expression changes associated with these inhibitors, indicative of heterogenicity in drug response (**Fig. S9**). Finally, we compared gene expression changes that are unique to RMC-6236 compared to trametinib. We saw a small number of genes that are up-and down-regulated by RMC-6236 in MEKi-sensitive and-resistant cells with little overlap between these groups (**Fig. 5F**). Taken together, these results indicate that RAS and MEK inhibition elicit an overlapping set of gene expression changes yet produce heterogeneous responses across individual cell lines with respect to growth inhibition.

**Figure 5.**
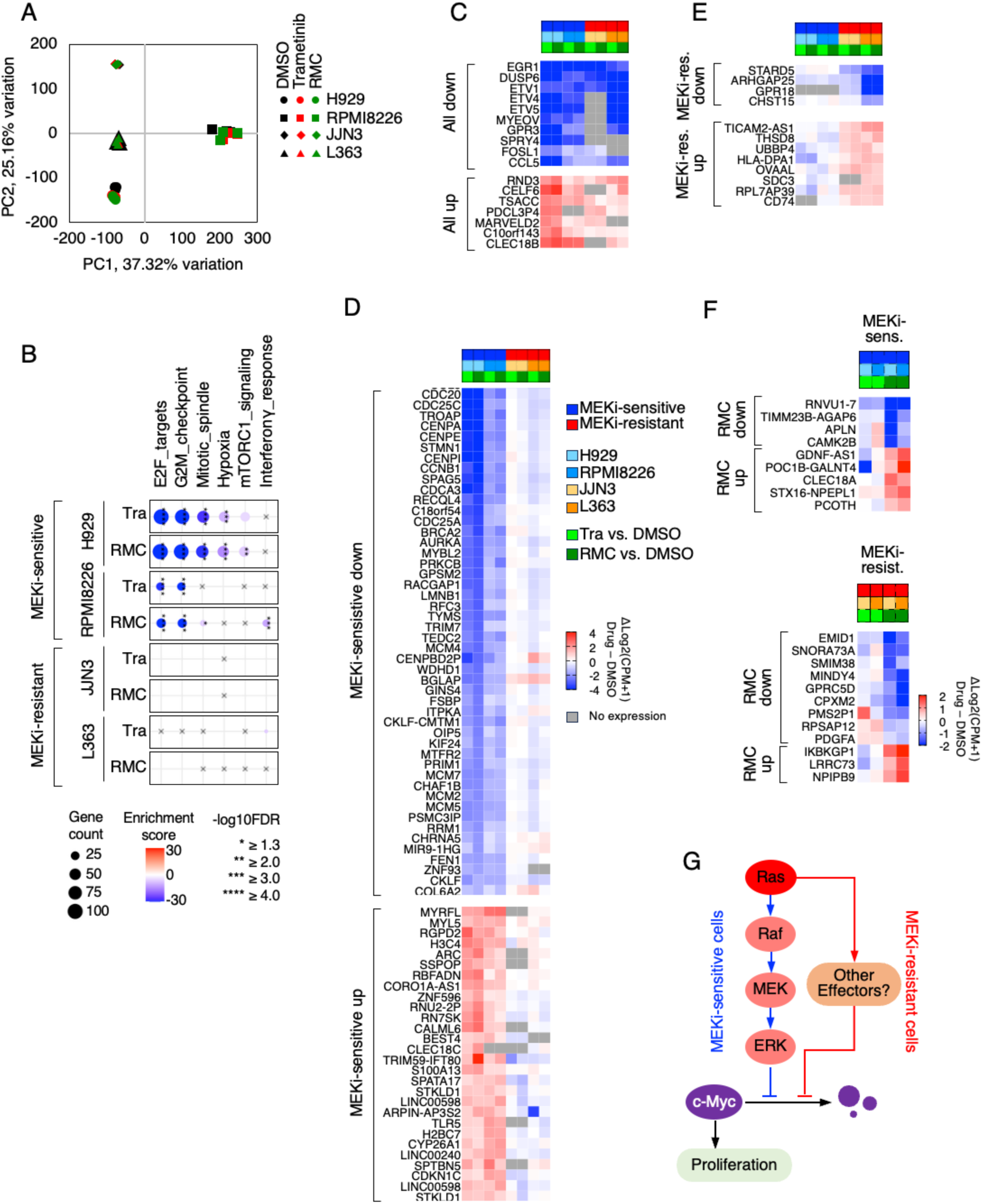
RASi and MEKi elicit overlapping transcriptional responses in MM cells. A. Principle component analysis (PCA) plot of RNAseq samples. MEKi-sensitive H929 and RPMI8226 cells and MEKi-resistant JJN3 and L363 cells were treated with trametinib (0.1 µM) or RMC-6236 (0.1 µM) for 4 days. Independent replicates were collected for RNA sequencing. B. Gene set enrichment analysis (GSEA) of Hallmark pathways altered by MEKi and RASi. C. Common gene expression changes induced by MEKi and RASi in all 4 cell lines. D. Gene expression changes induced by MEKi and RASi in in MEKi-sensitive H929 and RPMI8226 cells, but not in MEKi-resistant JJN3 and L363 cells. E. Gene expression changes induced by MEKi and RASi in MEKi-resistant JJN3 and L363 cells, but not in MEKi-sensitive H929 and RPMI8226 cells. F. Gene expression changes selectively induced by RMC-6236, but not by trametinib, in MEKi-sensitive H929 and RPMI8226 cells (top) and in MEKi-resistant JJN3 and L363 cell lines (bottom). G. A proposed model of how c-Myc stability is regulated by The Ras/MAPK pathway in Ras mutant MM cells.

## DISCUSSION

MM remains a challenging malignancy without targeted therapies. Frequent oncogenic mutation in *NRAS* and *KRAS* in MM, particularly in advanced and drug-refractory settings, implicates the Ras/MAPK pathway as a therapeutic target in this disease. Early clinical trials with MEK inhibitors, however, was met with limited single-agent activity in Ras mutant MM (20, 21), therefore raising the question of whether the Ras/MAPK pathway drives oncogene addiction in MM cells. In this study, we directly addressed this question by investigating the mechanism underlying the sensitivity of Ras mutant MM cells to the multi-Ras(ON) RASi RMC-6236 and to the MEKi trametinib. This study revealed several important insights. First, Ras-mutant MM cells remain addicted to the Ras oncogene for proliferation, as evidenced by their sensitivity to shRNA mediated Ras oncogene knockdown and to pharmacological inhibition of the Ras oncoprotein by RMC-6236. Second, Ras-mutant MM cells exhibit variable sensitivity to MEK inhibition, with some cell lines exhibiting strong MEKi resistance. Third, c-Myc protein stability is a key determinant of MEKi sensitivity in Ras mutant MM cells, as over-expression and genetic inhibition of c-Myc can decrease and increase MEKi sensitivity, respectively. Fourth, RASi downregulates c-Myc in all MM cell lines tested and c-Myc overexpression is insufficient to confer RASi resistance. Together, these results provide a molecular mechanism for the variable response to MEKi among Ras mutant MM cell lines and offers a potential explanation for the limited response of Ras mutant MM to MEKi that has been seen in clinical trials (20, 21). Importantly, our study provides strong pre-clinical evidence for targeting Ras directly in Ras mutant MM patients with multi-Ras(ON) inhibitors.

In MM pathogenesis, c-Myc is a frequent target of IgH translocations or secondary amplification, promoting MM cell proliferation and survival (48, 49). Because c-Myc expression is critical for the viability of MM cells (PMID: 35086951, 38714690 37934799). c-Myc has been actively studied as a therapeutic target across multiple cancer type (50, 51), including myeloma (52, 53). In solid tumors, c-Myc expression has been linked to resistance to inhibitors targeting the Ras/MAPK pathway (31, 54, 55). Our results extended these findings in the context of MM by showing c-Myc counteracts MAPK pathway dependence in MM cells, too. Although changes in several biomarkers correlated with MEKi sensitivity, including phospho-S6, cyclin D2 and c-Myc, only c-Myc stabilization contributed to MEKi resistance. Our results also revealed additional complexities in the relationship between Ras, MAPK, and c-Myc in MM cells that was not previously appreciated. First, in Ras mutant MM cells, ERK may not be the only kinase that phosphorylates S62 on c-Myc (11) as S62 phosphorylation persisted long after ERK kinase inactivation. Second, we found that although c-Myc level is insensitive to MEK inhibition in MEKi-resistant cells, it is still down-regulated by RASi. This suggests a model where c-Myc stability is regulated by MAPK in MEKi-sensitive cells, but by MAPK-independent Ras effectors in MEKi-resistant cells (**Fig. 5G**). Recent work showed that KRAS directly drives mTOR activation in MM cells (35), thus one possible mechanism of MAPK-independent c-Myc regulation by Ras could be through mTOR-driven c-Myc translation. Although inhibition of mTORC1 did not sensitize MEKi-resistant cells, further studies will be necessary to understand how the Ras/mTOR axis drives c-Myc over-expression in MM cells. Alternatively, it is plausible that c-Myc stability is differentially regulated by a E3 ligase, for example FBXW7, in MEKi-sensitive and-resistant cells, and future experiments will be necessary to identify such mechanism. Our finding provides further support to target c-Myc in MM as a therapeutic strategy (52).

Consistent with prior work describing a set of core MAPK pathway-regulated genes (45), we found these genes to be downregulated by both trametinib and RMC-6236 in MM cells. Both inhibitors drove a highly overlapping set of gene expression changes in each cell line, indicating the MAPK pathway to likely mediate a significant part of gene expression change downstream of mutant Ras. Trametinib elicited a larger gene expression change in MEKi-sensitive cells than in MEKi-resistant cells, consistent with a previous study demonstrating that a MAPK gene expression signature is only activated in a subset of Ras mutant MM cells (22). The small number of genes that are differentially regulated by RMC-6236 and trametinib in MEKi-resistant cells, however, suggest that RASi sensitivity in these cells might be associated with protein activity or stability, as in the case of c-Myc, rather than distinct gene expression alterations.

There are several limitations to our study. First, our study used a small panel of cell lines to model heterogeneity in Ras-mutant MM. INA6 cells have been reported to be IL-6 dependent (56). Our INA6 cells were authenticated and can be cultured without IL-6, likely reflecting a subline difference. Further experiment would be needed to test whether IL-6 can modulate Ras dependence in INA6 cells. Previous study has shown that JJN3 cells are insensitive to NRAS knockout (35). Our JJN3 cells were authenticated, and its NRAS dependence could reflect a subline variation. Testing our findings in a larger panel of Ras mutant MM cell lines will further generalize our conclusion. The Multi-Ras(ON) inhibitor RMC-6236 displays different degrees of potency towards different Ras alleles (25); thus, a larger panel of Ras MM cell line will further clarify its activity towards non-G12 mutant Ras alleles. As RMC-6236 can also block WT Ras signaling, it would be important to characterize its activity in Ras WT MM cells and in normal plasma cells to further confirm its selective toxicity in Ras mutant MM cells. Second, our study focused on the molecular dissection of drug sensitivity in vitro. Further studies in mouse models are needed to confirm our findings in vivo by testing the relative sensitive of xenograft tumors derived from MM cells to trametinib and to RMC-6236. Third, the mechanism by which Ras regulates c-Myc stability in MEKi-resistant MM cells is currently unclear, and further work is required to elucidate the mechanism. Fourth, our results showed that manipulating c-Myc protein alone is only able to partially alter MEKi sensitivity, suggesting contributions from other pathways that can cooperate with c-Myc to confer MEKi resistance. Last, our findings suggest that targeting the Ras/MAPK pathway in MM cells could lead to mostly cytostatic, rather than cytotoxic, responses. Identifying drug combinations that induce strong apoptosis in MM cells would therefore be a desirable future endeavor (30, 57).

RMC-6236 has demonstrated promising therapeutic benefit in pancreatic cancer and other solid tumors in recent clinical studies (25, 58). A potential translational implication of our study is that the new generation of multi-Ras(ON) inhibitors, as exemplified by RMC-6236, could provide better therapeutic benefit than MEKi in Ras mutant MM cells. These RASi can target multiple NRAS and KRAS mutations, offer a more consistent sensitivity profile among MM cell lines, and potentially have less on-target adverse effect. Importantly, c-Myc over-expression does not appear to overcome RASi sensitivity, thus reducing the potential for c-Myc mediated drug resistance. RMC-6236 has demonstrated promising therapeutic benefit in pancreatic cancer and other solid tumors in recent clinical studies (25, 58), our work provides a strong rationale for studies evaluating RMC-6236 and other multi-Ras(ON) inhibitors in the context of FDA-approved therapies for MM (17).

## MATERIALS AND METHODS

### Cell culture

Human MM cell lines (INA6, NCI-H929, RPMI8226, JJN3, L363, and KMS28BM) were obtained from Dr. Michael Kuehl. All cell lines were authenticated using Short Tandem Repeat analysis (LabCorp) within the last three years. MM cells were cultured in RPMI-1640 media (ThermoFisher Scientific Inc., cat. #SH30027FS) containing 10% fetal bovine serum (ThermoFisher Scientific Inc., cat. #10438026) and antibiotics (ThermoFisher Scientific Inc., cat. #15140163). Cells were maintained in a cell-culture incubator with humidified atmosphere with 5% CO_2_ and 95% air.

### Chemicals

Inhibitors and chemicals are obtained from the following commercial sources. From Selleck Chemicals: MEK1/2 inhibitor trametinib (cat. #S2673), doxorubicin (cat. #S1208), p70S6K inhibitor LY2584702 (cat. #S7704), GSK3β inhibitor CHIR99021 (cat. # S2924), AKT1/2/3 inhibitor MK-2206 2HCl (cat. # S1078), and proteasome inhibitor MG132 (cat. #S2619). From Chemgood: Ras(multi)(ON) inhibitor RMC-6236 (cat. #C-1418). From MedChemExpress: p38 inhibitor LY2228820 (cat. #HY-13241), CDK4/6 inhibitor palbociclib (cat. #HY-50767), JNK1/2/3 inhibitor JNK Inhibitor VIII (cat. #HY-107598), MEK5 inhibitor BIX02189 (cat. #HY-12056), and ERK5 inhibitor XMD8-92 (cat. #HY-14443). From Millipore Sigma: doxycycline (cat. #D9891), cycloheximide (cat. #C1988). From LC Laboratories: bortezomib (cat. #B-1408).

### Generation of lentiviral constructs

For expression vectors of human wild-type c-Myc (c-Myc-WT) and mutant c-Myc (c-Myc-T58A) with N-terminal 3×FLAG-tag were generated by PCR using EF1a_MYC_P2A_Hygro_Barcode (Addgene #120461) and EF1a_MYC T58A_P2A_Hygro_Barcode (Addgene #120513) as templates, respectively, using forward primer containing EcoRI restriction site (5’-CCGGAATTCAGATCTACCATGGACTACAAAGACCATGACGGTGATTATAAAGATCATGACATCGATTACAAGGATGACGATGACAAGGGAGGATCAGGAATGCCCCTCAACGTTAGCTTC-3’) and reverse primer containing BamHI restriction site (5’-CGCGGATCCTTACGCACAAGAGTTCCGTAG-3’) (Millipore Sigma). PCR amplicons and pLVX-puro destination vector were digested with EcoRI and BamHI and ligated at 16 °C overnight. The ligation products were transformed into *E. coli*, and plasmid DNA was isolated using a plasmid miniprep kit (QIAGEN, cat #27106). The desired clones were verified by whole plasmid sequencing (Quintara, MD, USA). For lentiviral vector expressing tet-inducible cyclin D2 with c-terminal HA-tag, human cyclin D2 cDNA was amplified by PCR using pRetroX-TRE3G-hCCND2 (kindly provided by Dr. Mardo Kõivomägi) as template with a forward primer containing SalI restriction site (5′-ACGCGTCGACGCCACCATGGAGCTGCTGTGCCACGAG-3′) and a reverse primer containing XhoI restriction site (5′-CCGCTCGAGTCAAGCGTAATCTGGAACATCGTATGGGTATCTGATCCTCCCAGGTCGATATCCCGCACGTC-3′). The pInducer-eGFP-HA destination vector was digested with SalI and XhoI to remove the eGFP cDNA fragment, and the cyclin D2 amplicon was inserted. Following bacterial transformation and plasmid purification, the desired clones were confirmed by whole-plasmid sequencing. Lentiviral vector expression tet-inducible omoMyc, pInducer-omomyc-HA, and control vector pInducer20-eGFP-HA were kindly gifted from Dr. William P. Tansey. Lentiviral shRNA vectors targeting human *KRAS* and *NRAS* from the MISSION® shRNA library (TRC1.0; Millipore Sigma) in the pLKO.1 backbone (shCtrl, GCCAAGATTCAGAATCCCAAA; shKRAS#1, GCAGACGTATATTGTATCATT; shKRAS#2, GAGGGCTTTCTTTGTGTATTT; shNRAS#1, GAAACCTGTTTGTTGGACATA; shNRAS#2, CAGTGCCATGAGAGACCAATA), as well as the second-generation lentiviral packaging plasmids psPAX2 (Addgene, cat. #12260) and pMD2.G (Addgene, cat. #12259), were kindly provided by Dr. Ryan M. Young.

### Generation of virally transduced stable cell lines

For viral vector packaging, 0.45 x 10^6^ HEK293FT cells were seeded into a 100 mm dish resulting in ∼70% confluency the next day. Lentiviral plasmid (6 µg) and packaging plasmids psPAX2 (4.5 µg) and pMD2.G (1.5 µg) were diluted in 1 mL of Opti-MEM™ (Gibco™, cat. # 31985070). Thirty-six µL of TransIT293 (Mirus Bio, cat. #MIR 2705) was then added to the diluted DNA and the mixture was incubated for 20 minutes at room temperature. Approximately 4 mL of media were removed from the dish, and 3 mL of fresh media was replenished. Transfection mixture was added dropwise to the HEK293FT cells. Virus-containing supernatants was collected daily up to 72 hours. After the last collection, virus was concentrated using Lenti-X™ Concentrator (Takara, cat. #631232) and the resulting virus pellet was resuspended using complete media to obtain high-titer virus stocks. For viral transduction, MM cells were counted and tested for cell viability. Cells were used for transduction only if viability is >80% by trypan blue staining (Gibco™, cat. #15250061). Cells were plated in a 6 well-plate at 3 x 10^6^ cell/well in 3 mL media. Polybrene (final concentration 10 µg/mL) and 200 µL of viral stock were added to each well. Spin infection was carried out by centrifugation for 90 minutes at 1,280 x g (2,500 rpm) using an Avanti J-15R centrifuge (Beckman Coulter). After centrifugation, the cells were recovered in an incubator for 2 days. For selection of stably transduced cells, cells were treated with puromycin (1 µg/mL) for 3 days.

### Cell cycle analysis

2 x 10⁶ MM cells were seeded onto a 6-well plate. The next day cells were treated with the inhibitors for 48 hours. Following treatment, cells were harvested and washed twice with phosphate-buffered saline (PBS), then resuspended in 300 µL of 1% FBS/PBS. While samples were continuously vortexed at low speed, 700 µL of ice-cold ethanol (100%) was added dropwise to the tube to minimize cell clumping. Cells were fixed for 1 hour at-20 °C, centrifuged, and washed twice with 1% FBS/PBS. Cell pellets were resuspended in 500 µL of buffer and transferred to 5 mL round-bottom FACS tube. Hoechst 33342 (final concentration 10 µg/mL; Biotium, cat. #40046) and RNase A (final concentration 100 µg/mL, Millipore Sigma, cat. #R6148) was added to all samples. Samples were incubated for 30 minutes at room temperature in the dark and analyzed on a Sony ID7000 Spectral Cell Analyzer (Sony Biotechnology Inc.) to quantify cell-cycle distribution and apoptotic populations.

### Western blot analysis

Cell lysates were prepared by lysing cells in cell lysis buffer (cell signaling technology, #9803). Protein concentrations were measured using Bradford protein assay dye reagent concentrate (Bio-Rad, cat. #5000006). Cell lysates were subjected to 8-16% sodium dodecyl sulfate polyacrylamide gel (Bio-Rad, cat. #5671104) electrophoresis followed by transfer to a nitrocellulose membrane (Bio-Rad, cat. #1704271). Membranes were blocked with blotting-grade milk (Bio-Rad, cat. #1706404) at 5%, incubated with primary antibodies listed in the table below diluted in Tris-buffered saline with 0.1% Tween-20 (TBST) at 4°C overnight, washed with TBST, and then incubated with horseradish peroxidase-conjugated secondary antibodies (Jackson ImmunoResearch, cat. #111-036-047 for rabbit and #115-036-072 for mouse, respectively). Protein signals were detected with a Western blotting detection reagent (ThermoFisher Scientific Inc., cat. #34095) in a ChemiDoc Imaging System (Bio-Rad, cat. #12003153). Densitometry quantification was carried out using Image Lab software (Bio-Rad, version 6.1).

| Target | Supplier | Cat. # |
| --- | --- | --- |
| KRAS | Millipore Sigma | WH0003845M1 |
| NRAS | Santa Cruz Biotechnology Inc. | sc-31 |
| p-p90RSK<sup>T359</sup> | Cell Signaling Technology, Inc. | 8753 |
| p90RSK | BD Biosciences | 610226 |
| p-p70S6K<sup>T389</sup> | Cell Signaling Technology, Inc. | 9234 |
| p70S6K | Cell Signaling Technology, Inc. | 9202 |
| p-S6<sup>S235/236</sup> | Cell Signaling Technology, Inc. | 4858 |
| S6 | Cell Signaling Technology, Inc. | 2217 |
| p-AKT<sup>S473</sup> | Cell Signaling Technology, Inc. | 4058 |
| AKT | Cell Signaling Technology, Inc. | 4691 |
| p-GSKβ<sup>S9</sup> | Cell Signaling Technology, Inc. | 5558 |
| GSKβ | Cell Signaling Technology, Inc. | 12456 |
| p-ERK1/2<sup>T202/Y204</sup> | Cell Signaling Technology, Inc. | 4377 |
| ERK1/2 | Cell Signaling Technology, Inc. | 9102 |
| DUSP4 | Cell Signaling Technology, Inc. | 5149 |
| DUSP6 | Cell Signaling Technology, Inc. | 39441 |
| Cyclin D1 | Millipore Sigma | C7464 |
| Cyclin D2 | Cell Signaling Technology, Inc. | 3741 |
| Cyclin D3 | Cell Signaling Technology, Inc. | 2936 |
| p-Myc<sup>T58</sup> | Cell Signaling Technology, Inc. | 46650 |
| p-Myc<sup>S62</sup> | Cell Signaling Technology, Inc. | 13748 |
| c-Myc | Cell Signaling Technology, Inc. | 18583 |
| PARP | Cell Signaling Technology, Inc. | 9542 |
| Cleaved-Casp3 | Cell Signaling Technology, Inc. | 9661 |
| Vinculin | Millipore Sigma | V9131 |
| α-Tubulin | Cell Signaling Technology, Inc. | 3873 |

### RT-qPCR

Total RNA was extracted using the RNeasy Mini QIAcube Kit (QIAGEN, cat. #74116). Reverse-transcribed cDNA was synthesized from purified RNA using the iScript™ gDNA Clear cDNA Synthesis Kit (Bio-Rad, cat. #1725035. qPCR reactions were performed on a CFX96™ Real-Time PCR Detection System (Bio-Rad). The primer sequences used for qPCR are as follows:

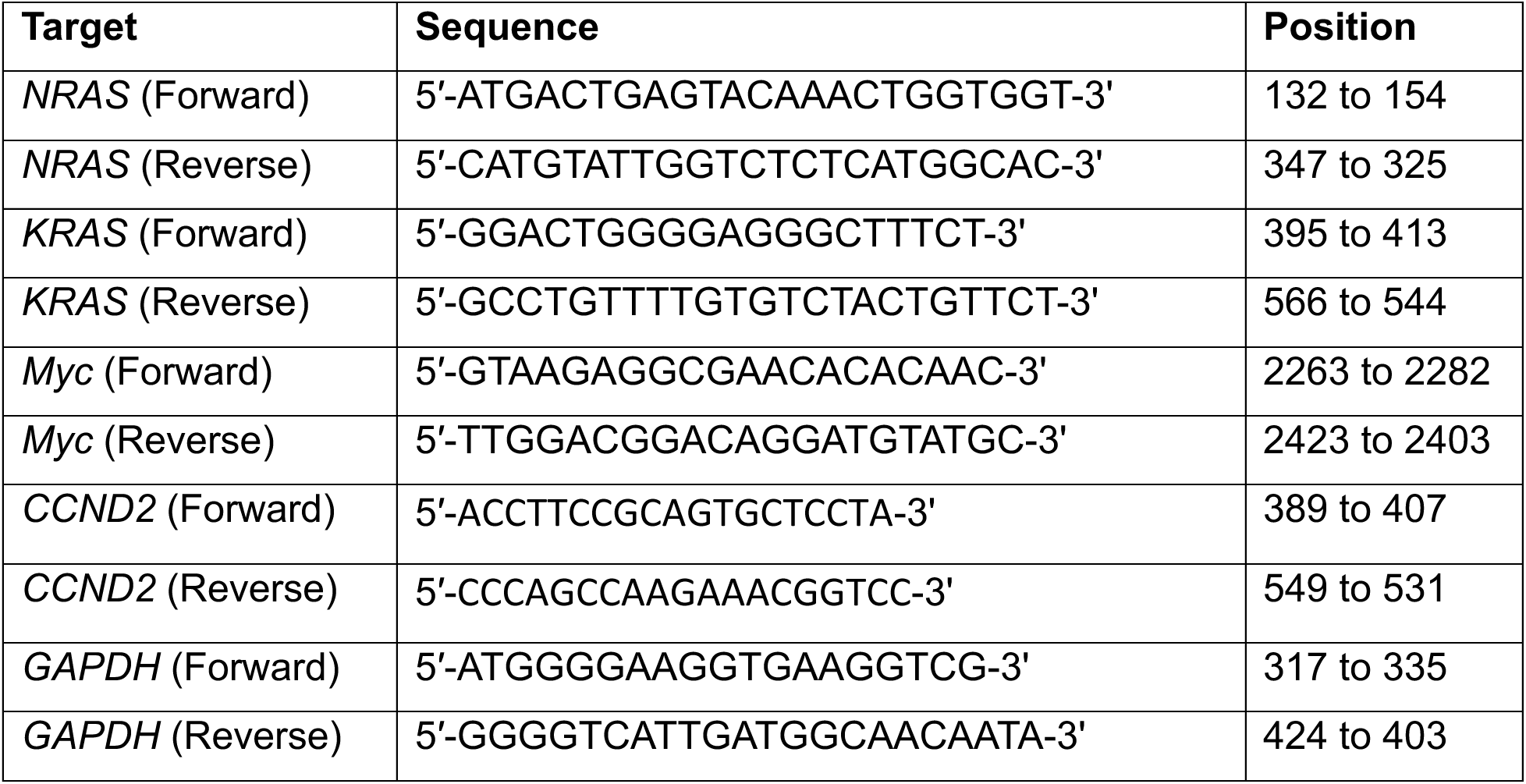

### Cell viability assay

For inhibitor treatment, cells were seeded in a 96-well plate at a density of 2,500/well. After one day, cells were treated with inhibitors for up to 4 days. After removing 100 µL of media, 100 µL of CellTiter-Glo solution (Promega, cat. #G7572) was added and the plate was incubated on shaker for 10 minutes at room temperature. Luminescence signals were read using a microplate reader (Tecan, Spark® Multimode Microplate Reader).

### Cycloheximide chase assay

MM cells were pretreated with vehicle or drug for 4 hours. Cells were then treated with cycloheximide (CHX; 20 µg/mL) to inhibit de novo protein synthesis. Cells were harvested at 0, 20, 40, and 60 minutes following CHX treatment. The proteasome inhibitor MG132 (10 µM) was added to a parallel 60-minute sample as a control for proteasome-mediated degradation. Cell lysates were subjected to Western blot analysis, and c-Myc protein levels were quantified by densitometry using vinculin as a loading control. The percentage of c-Myc protein remaining at each time point was calculated relative to the corresponding 0-minute sample. c-Myc protein half-life was estimated from the resulting decay curves using GraphPad Prism software. For statistical analysis, degradation-rate constants, *k*, were estimated separately from each independent biological replicate, and matched DMSO-and trametinib-treated samples were compared using a paired *t*-test on log-transformed *k* values.

### May-Grünwald staining

Human MM cells were seeded onto poly-D-lysine–coated 6-well plates (Becton Dickinson, cat. #354413) and incubated at 37 °C for 24 hours to allow cell attachment. After confirming adequate adherence, the culture medium was gently aspirated, and cells were stained with May-Grünwald solution (Millipore Sigma, cat. #63590) for 5 minutes. Cells were washed twice with PBS and subsequently incubated with Giemsa solution (Millipore Sigma, cat. #GS500) diluted 1:20 in distilled water for 20 minutes. Following staining, plates were rinsed briefly with water, air-dried, and imaged using a bright-field microscope.

### Bulk RNA-seq analysis

MM cells were treated with the indicated inhibitors for 4 days prior to collection. Each experimental condition was analyzed using three independent biological replicates. Total RNA was isolated using the RNeasy Mini QIAcube Kit (QIAGEN, cat. #74116) on a QIAcube Connect automated workstation (QIAGEN, cat. #9002864) according to the manufacturer’s instructions. RNA concentration was measured using the Quant-iT RiboGreen RNA Assay Kit (Thermo Fisher Scientific, cat. #R11490), and RNA integrity was assessed using the Agilent 4200 TapeStation system with RNA ScreenTape reagents (Agilent Technologies). Samples meeting quality-control criteria (RNA Integrity Number [RIN] ≥ 7) were advanced for library preparation. Strand-specific mRNA sequencing libraries were prepared by Psomagen Inc. (Rockville, MD, USA) using the Illumina Stranded mRNA Prep, Ligation Kit (Illumina, cat. #20040534). Briefly, 100 ng of total RNA was used as input for poly(A)-selected mRNA enrichment using oligo(dT) magnetic beads. Purified mRNA was fragmented and reverse-transcribed into first-strand cDNA using random hexamers in the presence of Actinomycin D to prevent DNA-dependent synthesis. Second-strand cDNA synthesis incorporated dUTP to preserve strand specificity. After end repair, A-tailing, and adapter ligation, libraries were amplified by PCR (13 cycles) using Illumina RNA UD Indexes (Set B; Illumina, cat. #20091657). Library quality and fragment-size distributions were evaluated using the Agilent 4200 TapeStation system with D5000 ScreenTape (Agilent Technologies, cat. #5067-5588), and libraries were quantified using the Quant-iT PicoGreen dsDNA Assay Kit (Thermo Fisher Scientific, cat. #P7589). Indexed libraries were normalized, pooled, and sequenced on an Illumina NovaSeq X Plus platform using the NovaSeq X Series 25B Reagent Kit (300-cycle; Illumina, cat. #20104706) to generate paired-end 150-bp reads. Base calling and demultiplexing were performed using Illumina software, and FASTQ files were generated using bcl2fastq (v2.20.0.422). For downstream RNA-seq analyses, raw RNA-seq counts generated by RSEM were analyzed using the iDEP web platform (version 2.01). Genes with counts per million (CPM) > 1 in three biological samples were retained for downstream analysis. For clustering and principal component analysis, count data were transformed using the edgeR log2(CPM + 1) method with a pseudocount of 1. Differential gene expression analysis was performed using the limma-voom pipeline, based on log₂ fold-change (FC), calculated as log₂(CPM+1) Drug-DMSO, with FDR < 0.05 and |log₂FC| ≥ 1 as cutoffs. Differentially expressed genes were subsequently used for heatmap visualization and Gene Set Enrichment Analysis (GSEA). Data were visualized using Excel, R, and GraphPad Prism. Raw sequencing data are deposited in GEO.

### Statistical analyses

All statistical analyses were carried out using GraphPad Prism software (version 10.6.1). Comparison of the means between two groups was determined by the Student’s *t*-test. Differences in the means among more than two groups were assessed by one-way ANOVA with Tukey’s multiple comparisons test. Unless otherwise specified, data are presented as mean ± standard error of the mean (SEM) and considered statistically significant if *p* < 0.05 (* *p* < 0.05, ** *p* < 0.01, *** *p* < 0.001, **** *p* < 0.0001).

## ACKNOWLEDGEMENTS

We thank Doug Lowy, Steven Cappell, Chih-Shia Lee, Vijaya Kumar Pidugu, and Wei-Chun Lee for advice; Marwa Afifi for reagents. The contributions of the NIH authors are considered works of the United States Government. The findings and conclusions presented in this paper are those of the authors and do not necessarily reflect the views of the NIH or the U.S. Department of Health and Human Services.

## AUTHOR CONTRIBUTIONS

**Y.-H.L.:** Conceptualization, methodology, investigation, formal analysis, visualization, original draft preparation, manuscript review and editing; **C.C.:** Methodology, investigation, manuscript review and editing; **H.Z.:** Methodology; **S.G.:** Methodology, resources, and investigation; **W.D.d.B.:** Methodology and investigation; **A.M.M.:** Bioinformatic analysis and data curation; **H.H.Y.:** Bioinformatic analysis and data curation; **T.J.M.:** Bioinformatic analysis and data curation; **R.M.Y.:** Conceptualization, resources, manuscript review and editing; **B.A.M.:** Conceptualization, resources, manuscript review and editing; **J.L.:** Conceptualization, supervision, project administration, funding acquisition, manuscript review and editing. All authors read and approved the final manuscript.

## SUPPLEMENTARY FIGURE LEGENDS

**Figure S1. Genetic and phenotypic characterization of the Ras-mutant MM cells**

A. Genetic alternations of the 6 MM cell lines used in this study. Hotspot mutations in commonly mutated genes, CRISPR KO gene dependency score (negative values indicate sgRNA lethality), and gene expression data were obtained from DepMap (https://depmap.org/portal/). Chromosome translocation (Tr) information was from the Keats lab portal (https://www.keatslab.org/myeloma-cell-lines/hmcl-characteristics).

B. Growth rates and MEKi and RASi dose-response parameters of MM cells. Baseline doubling time was calculated using the growth curves in panel S1C. IC_50_ was calculated as inhibitor concentration that produced a half-maximal response. Area under the curve (AUC) was calculated by integrating the entire dose-response curve as shown in Fig. 1C and 1D.

C. Baseline growth curves of MM cell lines. Cells were seeded at 2,500 cells/well in a 96-well plate and cell number was counted daily for 5 days.

D. Correlation plot of baseline doubling times and trametinib cell viability AUCs of MM cells. A lack of significant correlation between these two parameters is seen.

E. Sensitivity of Ras WT, NRAS and KRAS mutant MM cells to trametinib. Trametinib IC_50_ values of MM cell lines were plotted based on their Ras mutation status (30).

F. Correlation plot of Ras dependency and MEKi sensitivity in MM cells. Trametinib IC_50_ values of Ras mutant MM cell lines from (30) were plotted against NRAS or KRAS CRISPR scores from DepMap (https://depmap.org/portal/).

G. Sensitivity of MM cells to proteasome inhibition. Cells were treated with the proteasome inhibitor bortezomib (Bort) for 4 days and cell viability was measured using CellTiter-Glo assay.

H. Sensitivity of MM cells to chemotherapeutic agents. Cells were treated with the chemotherapy agent doxorubicin (Doxo) for 4 days and cell viability was measured using CellTiter-Glo assay.

**Figure S2. Trametinib causes growth arrest in MEKi-sensitive cell lines**

A. Effect of MEKi on cell cycle profile. Cells were treated with trametinib (Tra) for 48 hours, fixed and stained with Hoechst 33342, and cell cycle profile was analyzed using fluorescence-activated cell sorting (FACS).

B. Effect of MEKi on cell morphology. Cells were treated with trametinib for 48 hours then analyzed with May-Grünwald-Giemsa staining.

C. Effect of MEKi on apoptosis induction. Cells were treated with trametinib (Tra) or doxorubicin (Doxo) for 48 hours followed by immunoblotting of cleaved PARP and caspase 3 proteins.

**Figure S3. MEK inhibitor sensitivity is associated with downregulation of c-Myc, D-type cyclins and S6 phosphorylation**

A. Changes in protein analyte levels in response to MEKi. Cells were treated with trametinib for 4 days and cell lysates were immunoblotted for the indicated proteins and phospho-proteins. Representative blots from independent repeats are shown.

B. Protein analytes that are commonly downregulated by MEKi. Immunoblots from independent experiments shown in panel A were quantified by densitometry. Normalized proteins and phospho-proteins levels were plotted as dose-response curves against trametinib concentrations (graphs without error bars represent average of two independent experiments). Data from panels B, C and D are summarized as heatmap in Figure 2A.

C. Protein analytes that are downregulated by MEKi only in MEKi-sensitive cells (n=2 for p-p70S6K^T389^ in H929 and RPMI8226 cells).

D. Protein analytes that are not downregulated by MEKi.

E. Summary of correlation between cell viability and protein analyte changes induced by MEKi. Pair-wise Pearson correlation coefficient was calculated and plotted using AUC values of cell viability, protein and phospho-protein levels.

F. Correlation between cell viability and cyclin D2 and p-S6 changes induced by MEKi. AUC values are evaluated by Pearson correlation.

**Figure S4. RASi downregulates c-Myc, S6 phosphorylation, and cyclin D2**

A. Changes in protein analyte levels in response to RASi. Cells were treated with different doses of RMC-6236 (RMC) for 4 days and cell lysates were immunoblotted for the indicated proteins and phospho-proteins. Representative blots from independent repeats are shown.

B. Effect of RASi on p-S6 and cyclin D2 levels. Cells were treated with RMC-6236 for 4 days and pS6 and cyclin D2 levels was measured by immunoblotting and densitometry.

**Figure S5. Stability of c-Myc protein is differentially regulated in MEKi-sensitive and MEKi-resistant MM cells**

A. Effect of MEKi on c-Myc protein stability. Cells were treated with trametinib for 4 hours followed by cycloheximide (CHX) co-treatment for the indicated time to block protein translation. Co-treatment with the proteasome inhibitor MG132 was included as a control to block c-Myc degradation. Representative immunoblots from independent repeats quantified by densitometry and the data was plotted in Fig. 3B.

B. Effect of MEKi on c-Myc Ser62 phosphorylation. RPMI cells were treated with trametinib for the indicated time and changes in p-ERK, p-p90RSK, c-Myc phosphorylation at S62 (p-c-Myc^S62^) and total c-Myc level was immunoblotted.

C. Effect of MEKi on c-Myc Ser62 phosphorylation. INA6 and H929 cells were treated with trametinib for the indicated time and changes in p-ERK, p-c-Myc^S62^ and total c-Myc level was immunoblotted.

D. Effect of GSK3â inhibition on c-Myc T58 phosphorylation. RPMI8226 and H929 cells were treated with the GSK3β inhibitor CHIR99021 (CHIR) for 4 days and c-Myc phosphorylation at Thr58 (p-c-Myc^T58^) and total c-Myc protein levels was immunoblotted.

E. Effect of GSK3β inhibition on c-Myc protein level. Quantification of c-Myc protein levels in RPMI8226 and H929 cells were treated with trametinib (Tra) and CHIR99021 (CHRI) treatment as shown in Fig. 4A. Immunoblots from independent experiments were quantified by densitometry.

**Figure S6. S6K and AKT activity do not account for MEKi resistance**

A. Effect of p70S6K inhibition on S6 protein phosphorylation. Cells were treated with the selective p70S6K kinase inhibitor LY2584702 at indicated concentrations for 4 days and p-S6 level was immunoblotted.

B. Effect of p70S6K inhibition on MEKi sensitivity. MEKi-resistant JJN3, L363, and KMS28BM cells were treated with trametinib in combination with various doses of the p70S6K inhibitor LY2584702 for 4 days and cell viability was assessed using CellTiter-Glo assay.

C. Effect of mTORC1 inhibition on MEKi sensitivity. MEKi-resistant JJN3, L363, and KMS28BM cells were treated with trametinib in combination with various doses of the mTORC1 inhibitor rapamycin for 4 days and cell viability was assessed using CellTiter-Glo assay (n=1).

D. Effect of AKT inhibition on MEKi sensitivity. MEKi-resistant JJN3, L363, and KMS28BM cells were treated with trametinib in combination with various doses of the AKT inhibitor MK-2206 for 4 days and cell viability was assessed using CellTiter-Glo assay (n=2).

**Figure S7. Cyclin D2 protein and CDK4/6 activity do not account for MEKi resistance**

A. Effect of MEKi on cyclin D2 transcript level. Cells were treated with trametinib for 4 days and *CCND2* mRNA level was quantified by RT-qPCR.

B. Over-expression of exogenous cyclin D2. H929 cells were stably transduced with lentiviral vectors expressing tetracycline-inducible (tet-on) HA-tagged cDNAs of eGFP or cyclin D2. Cylin D2 expression was tested at indicated doses of doxycycline (Dox).

C. Effect of MEKi on cyclin D2 level. H929 cells inducibility expressing HA-tagged cyclin D2 was treated with trametinib and doxycycline either alone or in combination. Endogenous and exogenous Cylin D2 level was assessed by immunoblotting.

D. Effect of cyclin D2 overexpression on MEKi sensitivity. MEKi-sensitive H929 cells with (+Dox) and without (-Dox) eGFP or cyclin D2 overexpression were treated with trametinib for 4 days and cell viability was assessed using CellTiter-Glo assay.

E. Effect of CDK4/6 inhibition on MEKi sensitivity. MEKi-resistant JJN3, L363, and KMS28BM cells were treated with trametinib in combination with various doses of the CDK4/6 inhibitor palbociclib for 4 days and cell viability was assessed using CellTiter-Glo assay.

**Figure S8. Alternative MAPK pathways do not account for MEKi resistance** A-D. Effect of MAPK inhibitors on MEKi sensitivity. MEKi-resistant JJN3, L363, and KMS28BM cells were treated with trametinib in combination with various doses of the MEK5 kinase inhibitor BIX02189 (A), ERK5 kinase inhibitor XMD8-92 (B), p38 kinase inhibitor LY2228820 (C), and the JNK kinase inhibitor JNK inhibitor VIII (D) for 4 days and cell viability was assessed using CellTiter-Glo assay (n=1).

**Figure S9. Cell-line specific gene expression changes induced by MEKi and RASi**

Heatmaps showing genes that are up-and down-regulated by both trametinib and RMC-6236 that are specific to each individual cell line.

## REFERENCES

1. Siegel RL, Kratzer TB, Giaquinto AN, Sung H, Jemal A. Cancer statistics, 2025. CA Cancer J Clin. 2025;75(1):10–45.

2. Chesi M, Nardini E, Lim RS, Smith KD, Kuehl WM, Bergsagel PL. The t(4;14) translocation in myeloma dysregulates both FGFR3 and a novel gene, MMSET, resulting in IgH/MMSET hybrid transcripts. Blood. 1998;92(9):3025–34.

3. Mahindra A, Hideshima T, Anderson KC. Multiple myeloma: biology of the disease. Blood Rev. 2010;24 Suppl 1:S5–11.

4. Mikulasova A, Ashby C, Tytarenko RG, Qu P, Rosenthal A, Dent JA, et al. Microhomology-mediated end joining drives complex rearrangements and overexpression of MYC and PVT1 in multiple myeloma. Haematologica. 2020;105(4):1055–66.

5. Landgren O, Hofmann JN, McShane CM, Santo L, Hultcrantz M, Korde N, et al. Association of Immune Marker Changes With Progression of Monoclonal Gammopathy of Undetermined Significance to Multiple Myeloma. JAMA Oncol. 2019;5(9):1293–301.

6. Landgren O, Kyle RA, Pfeiffer RM, Katzmann JA, Caporaso NE, Hayes RB, et al. Monoclonal gammopathy of undetermined significance (MGUS) consistently precedes multiple myeloma: a prospective study. Blood. 2009;113(22):5412–7.

7. Weiss BM, Abadie J, Verma P, Howard RS, Kuehl WM. A monoclonal gammopathy precedes multiple myeloma in most patients. Blood. 2009;113(22):5418–22.

8. Schavgoulidze A, Corre J, Samur MK, Mazzotti C, Pavageau L, Perrot A, et al. RAS/RAF landscape in monoclonal plasma cell conditions. Blood. 2024;144(2):201–5.

9. Leone G, DeGregori J, Sears R, Jakoi L, Nevins JR. Myc and Ras collaborate in inducing accumulation of active cyclin E/Cdk2 and E2F. Nature. 1997;387(6631):422–6.

10. Albanese C, Johnson J, Watanabe G, Eklund N, Vu D, Arnold A, et al. Transforming p21ras mutants and c-Ets-2 activate the cyclin D1 promoter through distinguishable regions. J Biol Chem. 1995;270(40):23589–97.

11. Sears R, Nuckolls F, Haura E, Taya Y, Tamai K, Nevins JR. Multiple Ras-dependent phosphorylation pathways regulate Myc protein stability. Genes Dev. 2000;14(19):2501–14.

12. Sears R, Leone G, DeGregori J, Nevins JR. Ras enhances Myc protein stability. Mol Cell. 1999;3(2):169–79.

13. Soucek L, Whitfield J, Martins CP, Finch AJ, Murphy DJ, Sodir NM, et al. Modelling Myc inhibition as a cancer therapy. Nature. 2008;455(7213):679–83.

14. Neri P, Barwick BG, Jung D, Patton JC, Maity R, Tagoug I, et al. ETV4-Dependent Transcriptional Plasticity Maintains MYC Expression and Results in IMiD Resistance in Multiple Myeloma. Blood Cancer Discov. 2024;5(1):56–73.

15. Maura F, Coffey DG, Stein CK, Braggio E, Ziccheddu B, Sharik ME, et al. The genomic landscape of Vk*MYC myeloma highlights shared pathways of transformation between mice and humans. Nat Commun. 2024;15(1):3844.

16. Shi Y, Sun F, Cheng Y, Holmes B, Dhakal B, Gera JF, et al. Critical Role for Cap-Independent c-MYC Translation in Progression of Multiple Myeloma. Mol Cancer Ther. 2022;21(4):502–10.

17. Rajkumar SV. Multiple myeloma: 2024 update on diagnosis, risk-stratification, and management. Am J Hematol. 2024;99(9):1802–24.

18. National Cancer Institute. SEER Cancer Stat Facts: Myeloma: National Cancer Institute; 2026 [Available from: https://seer.cancer.gov/statfacts/html/mulmy.html.

19. Xu J, Pfarr N, Endris V, Mai EK, Md Hanafiah NH, Lehners N, et al. Molecular signaling in multiple myeloma: association of RAS/RAF mutations and MEK/ERK pathway activation. Oncogenesis. 2017;6(5):e337.

20. Holkova B, Zingone A, Kmieciak M, Bose P, Badros AZ, Voorhees PM, et al. A Phase II Trial of AZD6244 (Selumetinib, ARRY-142886), an Oral MEK1/2 Inhibitor, in Relapsed/Refractory Multiple Myeloma. Clin Cancer Res. 2016;22(5):1067–75.

21. Guo C, Chenard-Poirier M, Roda D, de Miguel M, Harris SJ, Candilejo IM, et al. Intermittent schedules of the oral RAF-MEK inhibitor CH5126766/VS-6766 in patients with RAS/RAF-mutant solid tumours and multiple myeloma: a single-centre, open-label, phase 1 dose-escalation and basket dose-expansion study. Lancet Oncol. 2020;21(11):1478–88.

22. Lin YT, Way GP, Barwick BG, Mariano MC, Marcoulis M, Ferguson ID, et al. Integrated phosphoproteomics and transcriptional classifiers reveal hidden RAS signaling dynamics in multiple myeloma. Blood Adv. 2019;3(21):3214–27.

23. Sriskandarajah P, De Haven Brandon A, MacLeod K, Carragher NO, Kirkin V, Kaiser M, et al. Combined targeting of MEK and the glucocorticoid receptor for the treatment of RAS-mutant multiple myeloma. BMC Cancer. 2020;20(1):269.

24. Riedl JM, Matsubara H, McNeil R, Patel PS, Fece de la Cruz F, Gulhan DC, et al. Emerging landscape of KRAS inhibitors in cancer treatment. Cancer Cell. 2026;44(3):471–97.

25. Jiang J, Jiang L, Maldonato BJ, Wang Y, Holderfield M, Aronchik I, et al. Translational and Therapeutic Evaluation of RAS-GTP Inhibition by RMC-6236 in RAS-Driven Cancers. Cancer Discov. 2024;14(6):994–1017.

26. O’Reilly EM, Wainberg ZA, Hendifar AE, Borad MJ, Pietrantonio F, Pant S, et al. Daraxonrasib or Chemotherapy in Previously Treated Metastatic Pancreatic Cancer. N Engl J Med. 2026.

27. Tsherniak A, Vazquez F, Montgomery PG, Weir BA, Kryukov G, Cowley GS, et al. Defining a Cancer Dependency Map. Cell. 2017;170(3):564–76 e16.

28. Delmer A, Ajchenbaum-Cymbalista F, Tang R, Ramond S, Faussat AM, Marie JP, et al. Overexpression of cyclin D2 in chronic B-cell malignancies. Blood. 1995;85(10):2870–6.

29. Suzuki R, Kuroda H, Komatsu H, Hosokawa Y, Kagami Y, Ogura M, et al. Selective usage of D-type cyclins in lymphoid malignancies. Leukemia. 1999;13(9):1335–42.

30. Hughitt VK, Simmons JK, Gorjifard S, Michalowski A, Wilson K, Zhang X, et al. Large-scale human myeloma cell line small molecule compound screen dataset. Sci Data. 2025;12(1):661.

31. Vaseva AV, Blake DR, Gilbert TSK, Ng S, Hostetter G, Azam SH, et al. KRAS Suppression-Induced Degradation of MYC Is Antagonized by a MEK5-ERK5 Compensatory Mechanism. Cancer Cell. 2018;34(5):807–22 e7.

32. Luscher B, Eisenman RN. Proteins encoded by the c-myc oncogene: analysis of c-myc protein degradation. Princess Takamatsu Symp. 1986;17:291–301.

33. Welcker M, Orian A, Jin J, Grim JE, Harper JW, Eisenman RN, et al. The Fbw7 tumor suppressor regulates glycogen synthase kinase 3 phosphorylation-dependent c-Myc protein degradation. Proc Natl Acad Sci U S A. 2004;101(24):9085–90.

34. Soucek L, Helmer-Citterich M, Sacco A, Jucker R, Cesareni G, Nasi S. Design and properties of a Myc derivative that efficiently homodimerizes. Oncogene. 1998;17(19):2463–72.

35. Yang Y, Bolomsky A, Oellerich T, Chen P, Ceribelli M, Haupl B, et al. Oncogenic RAS commandeers amino acid sensing machinery to aberrantly activate mTORC1 in multiple myeloma. Nat Commun. 2022;13(1):5469.

36. Tolcher A, Goldman J, Patnaik A, Papadopoulos KP, Westwood P, Kelly CS, et al. A phase I trial of LY2584702 tosylate, a p70 S6 kinase inhibitor, in patients with advanced solid tumours. Eur J Cancer. 2014;50(5):867–75.

37. Carracedo A, Ma L, Teruya-Feldstein J, Rojo F, Salmena L, Alimonti A, et al. Inhibition of mTORC1 leads to MAPK pathway activation through a PI3K-dependent feedback loop in human cancer. J Clin Invest. 2008;118(9):3065–74.

38. Lim KH, Counter CM. Reduction in the requirement of oncogenic Ras signaling to activation of PI3K/AKT pathway during tumor maintenance. Cancer Cell. 2005;8(5):381–92.

39. Dey A, She H, Kim L, Boruch A, Guris DL, Carlberg K, et al. Colony-stimulating factor-1 receptor utilizes multiple signaling pathways to induce cyclin D2 expression. Mol Biol Cell. 2000;11(11):3835–48.

40. Ando K, Ajchenbaum-Cymbalista F, Griffin JD. Regulation of G1/S transition by cyclins D2 and D3 in hematopoietic cells. Proc Natl Acad Sci U S A. 1993;90(20):9571–5.

41. Fry DW, Harvey PJ, Keller PR, Elliott WL, Meade M, Trachet E, et al. Specific inhibition of cyclin-dependent kinase 4/6 by PD 0332991 and associated antitumor activity in human tumor xenografts. Mol Cancer Ther. 2004;3(11):1427–38.

42. Tatake RJ, O’Neill MM, Kennedy CA, Wayne AL, Jakes S, Wu D, et al. Identification of pharmacological inhibitors of the MEK5/ERK5 pathway. Biochem Biophys Res Commun. 2008;377(1):120–5.

43. Patnaik A, Haluska P, Tolcher AW, Erlichman C, Papadopoulos KP, Lensing JL, et al. A First-in-Human Phase I Study of the Oral p38 MAPK Inhibitor, Ralimetinib (LY2228820 Dimesylate), in Patients with Advanced Cancer. Clin Cancer Res. 2016;22(5):1095–102.

44. Szczepankiewicz BG, Kosogof C, Nelson LT, Liu G, Liu B, Zhao H, et al. Aminopyridine-based c-Jun N-terminal kinase inhibitors with cellular activity and minimal cross-kinase activity. J Med Chem. 2006;49(12):3563–80.

45. Klomp JA, Klomp JE, Stalnecker CA, Bryant KL, Edwards AC, Drizyte-Miller K, et al. Defining the KRAS-and ERK-dependent transcriptome in KRAS-mutant cancers. Science. 2024;384(6700).

46. Villalonga P, Guasch RM, Riento K, Ridley AJ. RhoE inhibits cell cycle progression and Ras-induced transformation. Mol Cell Biol. 2004;24(18):7829–40.

47. Liu G, Zhang Q, Xia L, Shi M, Cai J, Zhang H, et al. RNA-binding protein CELF6 is cell cycle regulated and controls cancer cell proliferation by stabilizing p21. Cell Death Dis. 2019;10(10):688.

48. Avet-Loiseau H, Gerson F, Magrangeas F, Minvielle S, Harousseau JL, Bataille R, et al. Rearrangements of the c-myc oncogene are present in 15% of primary human multiple myeloma tumors. Blood. 2001;98(10):3082–6.

49. Walker BA, Wardell CP, Brioli A, Boyle E, Kaiser MF, Begum DB, et al. Translocations at 8q24 juxtapose MYC with genes that harbor superenhancers resulting in overexpression and poor prognosis in myeloma patients. Blood Cancer J. 2014;4(3):e191.

50. Duffy MJ, Tang M, Crown J. MYC as a Target for Cancer Treatment: from Undruggable to Druggable? Target Oncol. 2025;20(5):791–801.

51. Dang CV. MYC on the path to cancer. Cell. 2012;149(1):22–35.

52. Gaikwad SM, Phyo Z, Arteaga AQ, Gorjifard S, Calabrese DR, Connors D, et al. A Small Molecule Stabilizer of the MYC G4-Quadruplex Induces Endoplasmic Reticulum Stress, Senescence and Pyroptosis in Multiple Myeloma. Cancers (Basel). 2020;12(10).

53. Martinez-Martin S, Soucek L. MYC inhibitors in multiple myeloma. Cancer Drug Resist. 2021;4(4):842–65.

54. Zhao Y, Murciano-Goroff YR, Xue JY, Ang A, Lucas J, Mai TT, et al. Diverse alterations associated with resistance to KRAS(G12C) inhibition. Nature. 2021;599(7886):679–83.

55. Dilly J, Hoffman MT, Abbassi L, Li Z, Paradiso F, Parent BD, et al. Mechanisms of Resistance to Oncogenic KRAS Inhibition in Pancreatic Cancer. Cancer Discov. 2024;14(11):2135–61.

56. Burger R, Guenther A, Bakker F, Schmalzing M, Bernand S, Baum W, et al. Gp130 and ras mediated signaling in human plasma cell line INA-6: a cytokine-regulated tumor model for plasmacytoma. Hematol J. 2001;2(1):42–53.

57. Peat TJ, Gaikwad SM, Dubois W, Gyabaah-Kessie N, Zhang S, Gorjifard S, et al. Drug combinations identified by high-throughput screening promote cell cycle transition and upregulate Smad pathways in myeloma. Cancer Lett. 2023;568:216284.

58. Cregg J, Edwards AV, Chang S, Lee BJ, Knox JE, Tomlinson ACA, et al. Discovery of Daraxonrasib (RMC-6236), a Potent and Orally Bioavailable RAS(ON) Multi-selective, Noncovalent Tri-complex Inhibitor for the Treatment of Patients with Multiple RAS-Addicted Cancers. J Med Chem. 2025;68(6):6064–83.

