## Supplementary Figure File for "Stability of c-Myc protein differentiates Ras oncogene addiction and MAPK pathway dependency in Ras-mutant multiple myeloma"

### Figure S1

A

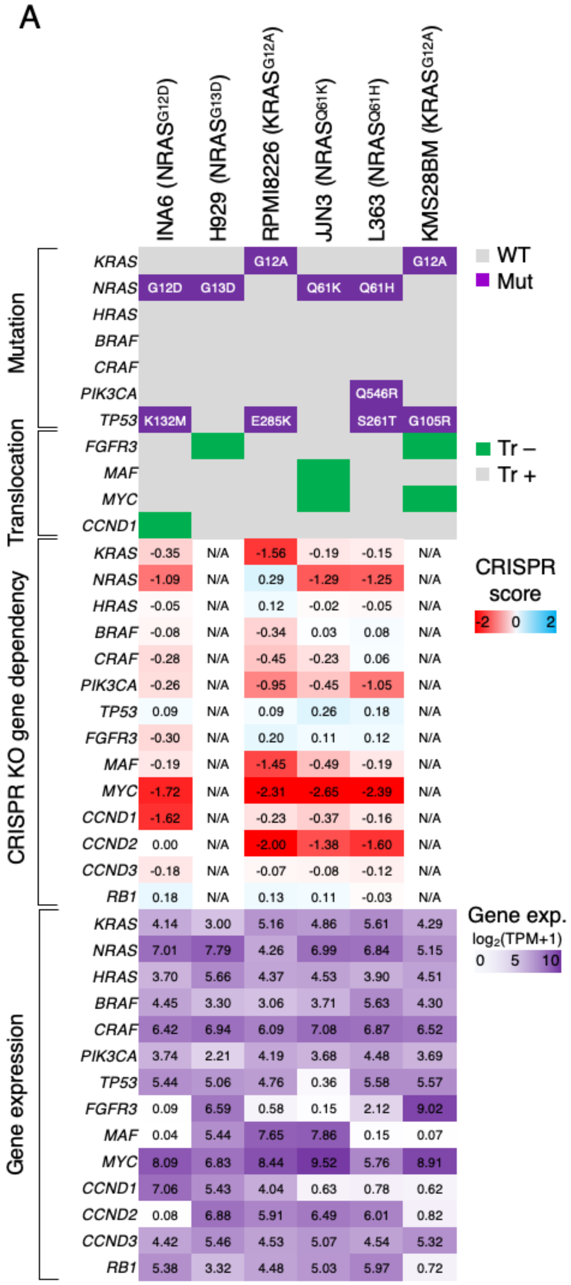


B

| Cell line | Doubling Time (h) | RMC-6236 dose-response Bottom | RMC-6236 dose-response IC50 (nM) | RMC-6236 dose-response AUC (a.u.) | Trametinib dose-response Bottom | Trametinib dose-response IC50 (nM) | Trametinib dose-response AUC (a.u.) |
|---|---|---|---|---|---|---|---|
| INA6 | 37.6 | 0.11 | 0.27 | 2.03 | 0.22 | 1.31 | 2.79 |
| H929 | 22.7 | 0.23 | 5.42 | 3.43 | 0.35 | 7.22 | 3.67 |
| RPMI8226 | 28.6 | 0.39 | 0.75 | 3.16 | 0.53 | 1.54 | 3.69 |
| JJN3 | 25.2 | 0.63 | 15.0 | 4.29 | 0.76 | 90.0 | 4.83 |
| L363 | 23.3 | 0.53 | 0.83 | 3.93 | 0.87 | 506 | 5.05 |
| KMS28BM | 30.8 | 0.57 | 2.23 | 3.93 | 0.88 | 1468 | 5.12 |

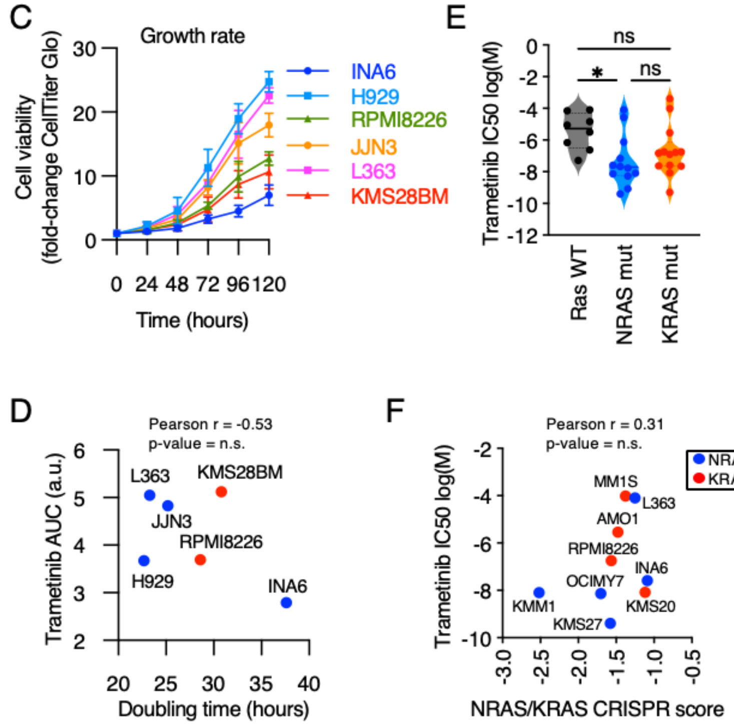

Figure S1 (cont.)

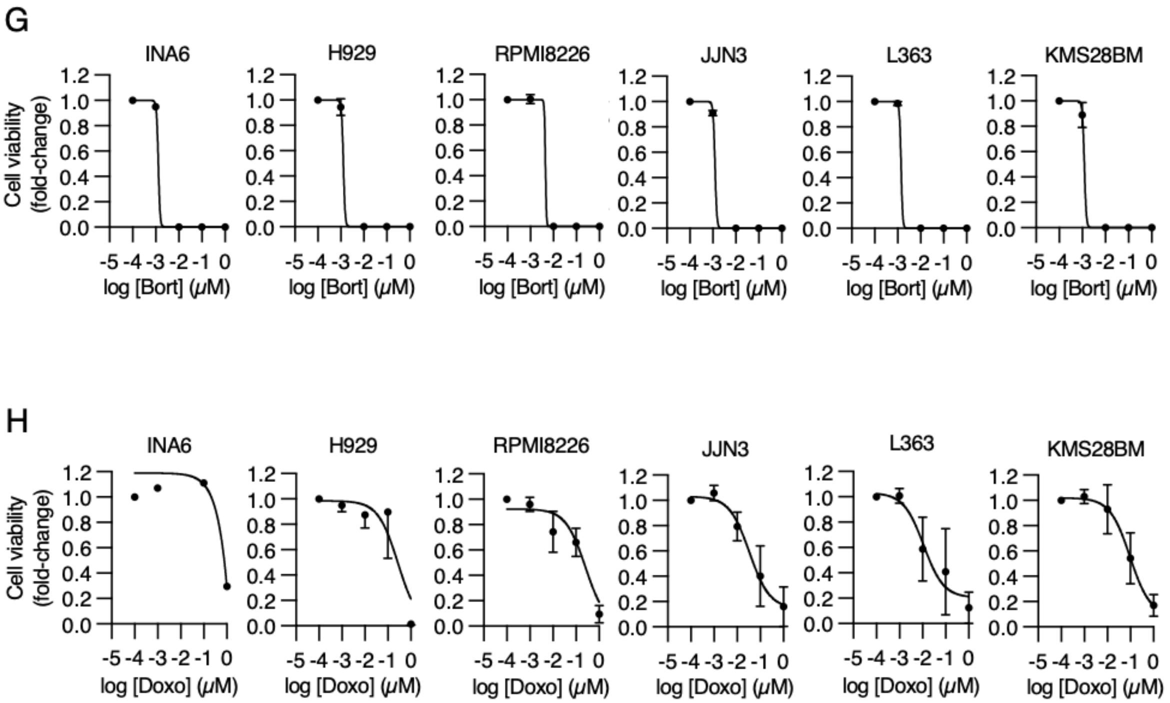

Figure S2

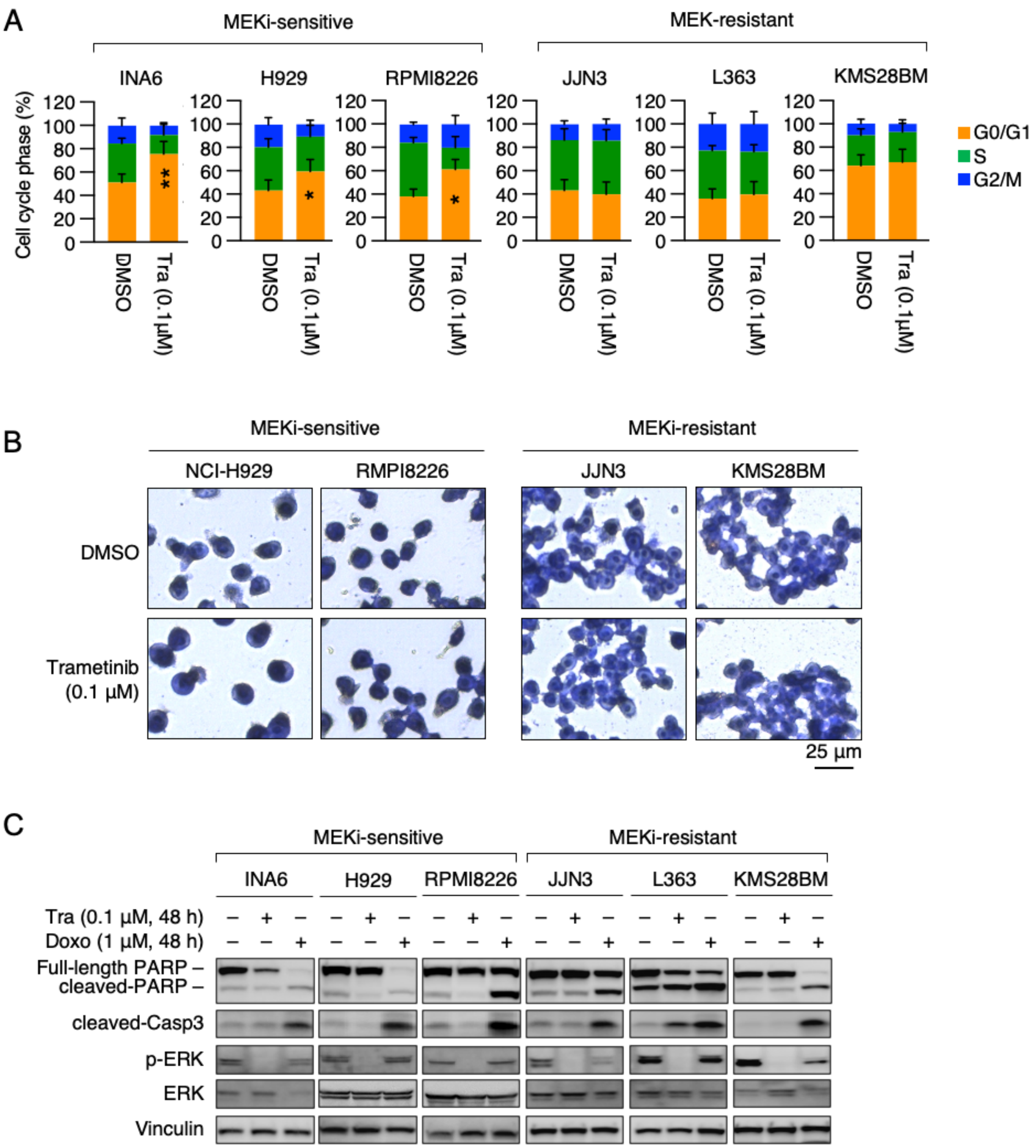

A
MEKi-sensitive
MEK-resistant
INA6
H929
RPMI8226
JJN3
L363
KMS28BM
Cell cycle phase (%)
DMSO
Tra (0.1µM)
G0/G1
S
G2/M
B
MEKi-sensitive
MEKi-resistant
NCI-H929
RMPI8226
JJN3
KMS28BM
DMSO
Trametinib (0.1 µM)
25 µm
C
MEKi-sensitive
MEKi-resistant
INA6
H929
RPMI8226
JJN3
L363
KMS28BM
Tra (0.1 µM, 48 h)
Doxo (1 µM, 48 h)
Full-length PARP
cleaved-PARP
cleaved-Casp3
p-ERK
ERK
Vinculin

### Figure S3

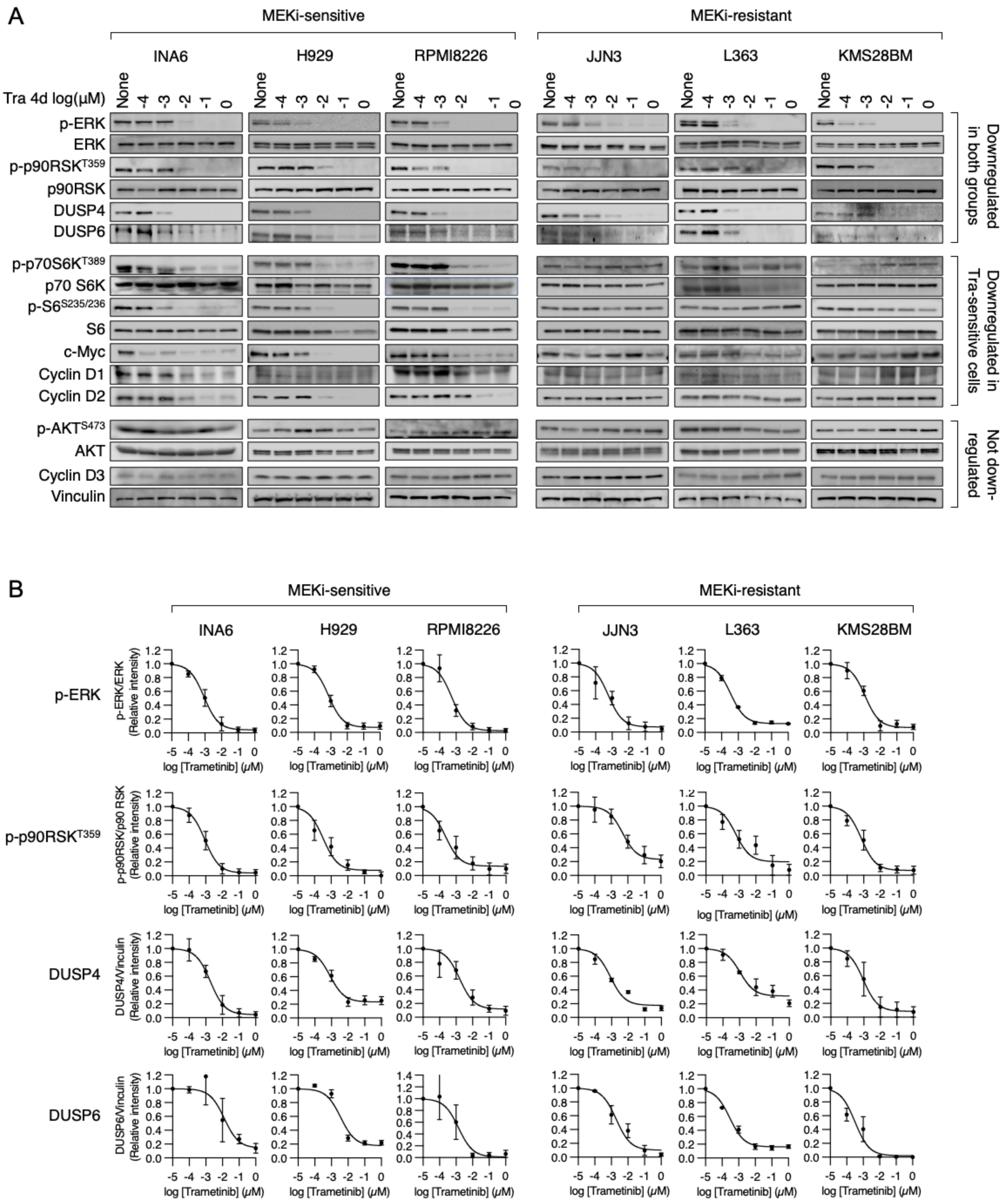

A
MEKi-sensitive
MEKi-resistant
INA6
H929
RPMI8226
JJN3
L363
KMS28BM
Tra 4d log(μM)
None
-4
-3
-2
-1
0
p-ERK
ERK
p-p90RSKT359
p90RSK
DUSP4
DUSP6
p-p70S6KT389
p70 S6K
p-S6S235/236
S6
c-Myc
Cyclin D1
Cyclin D2
p-AKTS473
AKT
Cyclin D3
Vinculin
Downregulated in both groups
Downregulated in Tra-sensitive cells
Not down-regulated
B
MEKi-sensitive
MEKi-resistant
INA6
H929
RPMI8226
JJN3
L363
KMS28BM
p-ERK
p-ERK/ERK (Relative intensity)
p-p90RSKT359
p-p90RSK/p90 RSK (Relative intensity)
DUSP4
DUSP4/Vinculin (Relative intensity)
DUSP6
DUSP6/Vinculin (Relative intensity)
log [Trametinib] (μM)

Figure S3 (cont.)

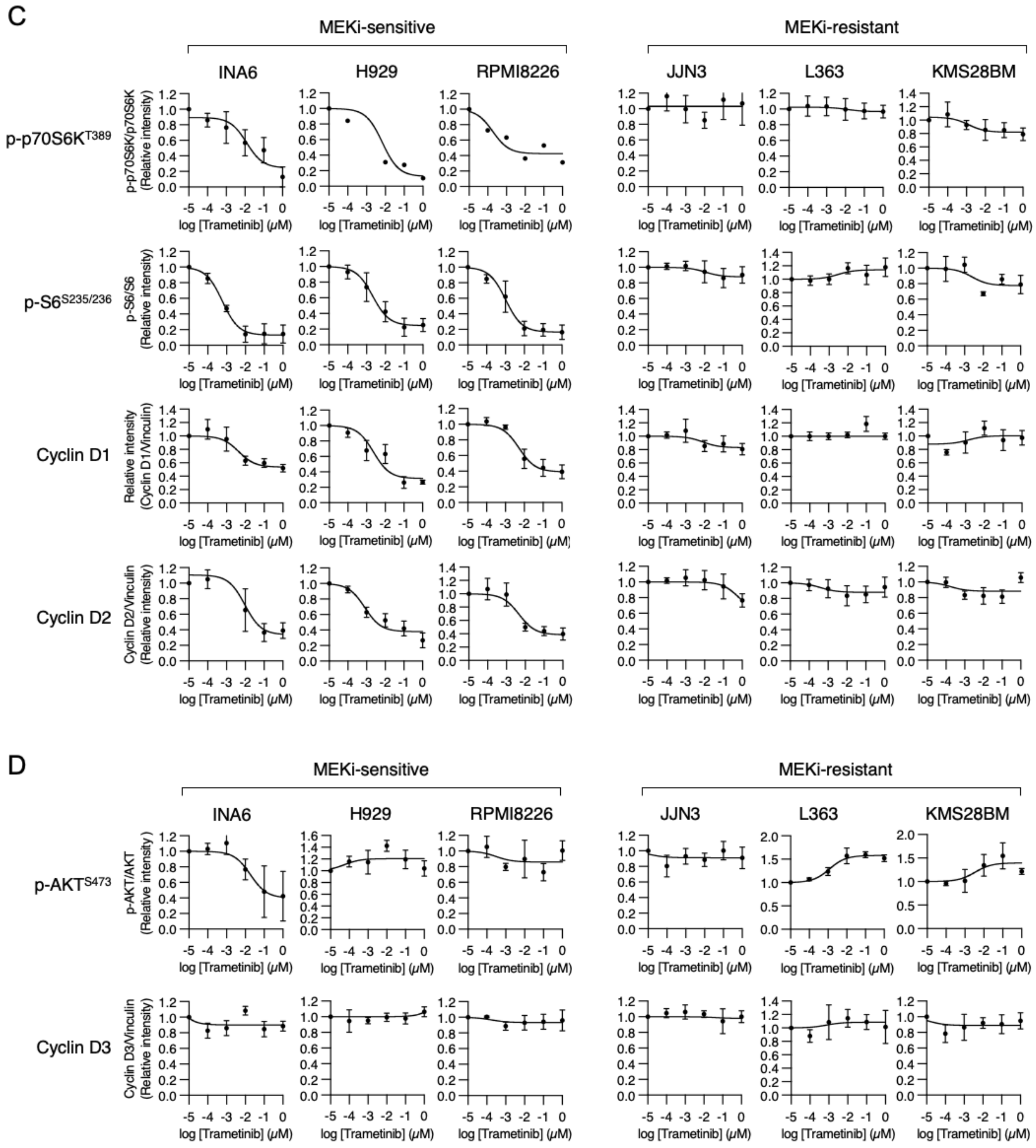

### Figure S3 (cont.)

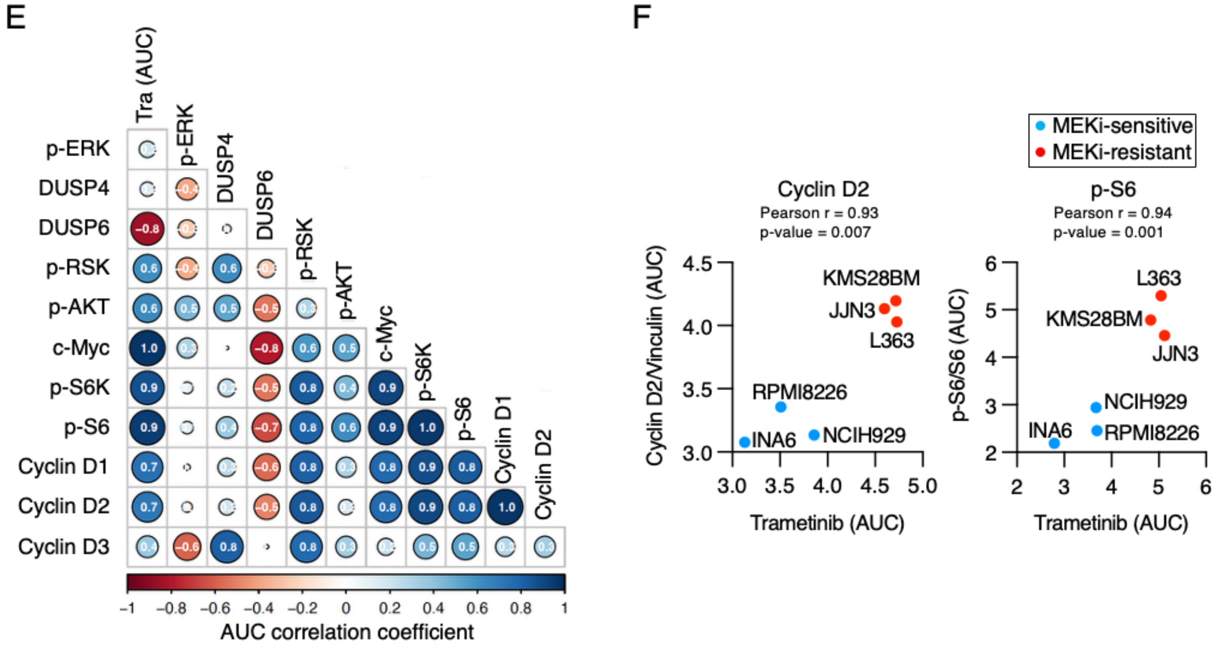

Figure S4.

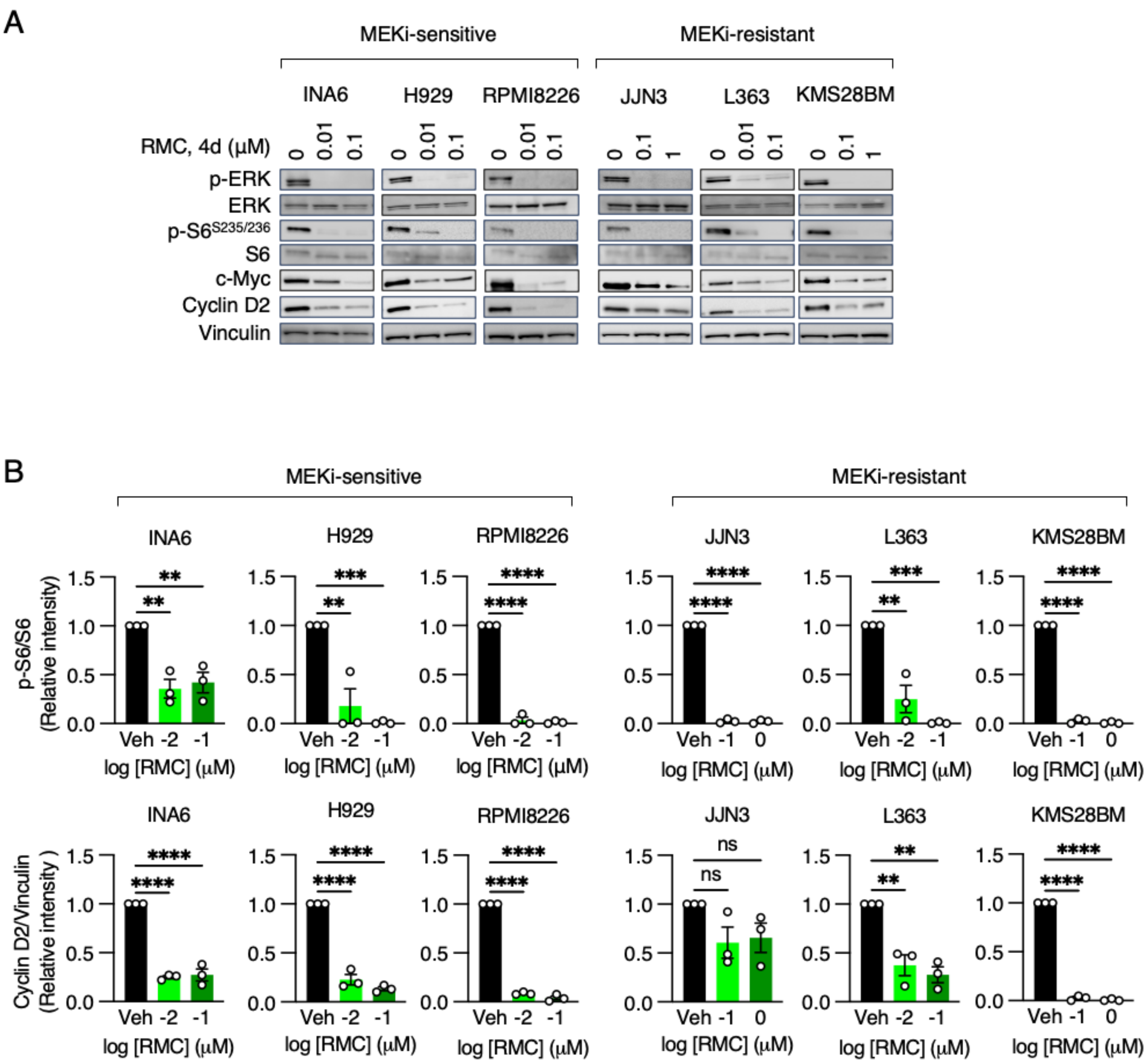
A
MEKi-sensitive
MEKi-resistant
INA6
H929
RPMI8226
JJN3
L363
KMS28BM
RMC, 4d (μM)
0 0.01 0.1
0 0.01 0.1
0 0.01 0.1
0 0.1 1
0 0.01 0.1
0 0.1 1
p-ERK
ERK
p-S6S235/236
S6
c-Myc
Cyclin D2
Vinculin
B
MEKi-sensitive
MEKi-resistant
p-S6/S6 (Relative intensity)
Cyclin D2/Vinculin (Relative intensity )
Veh -2 -1
Veh -1 0
log [RMC] (μM)
ns
1.5
1.0
0.5
0.0

### Figure S5

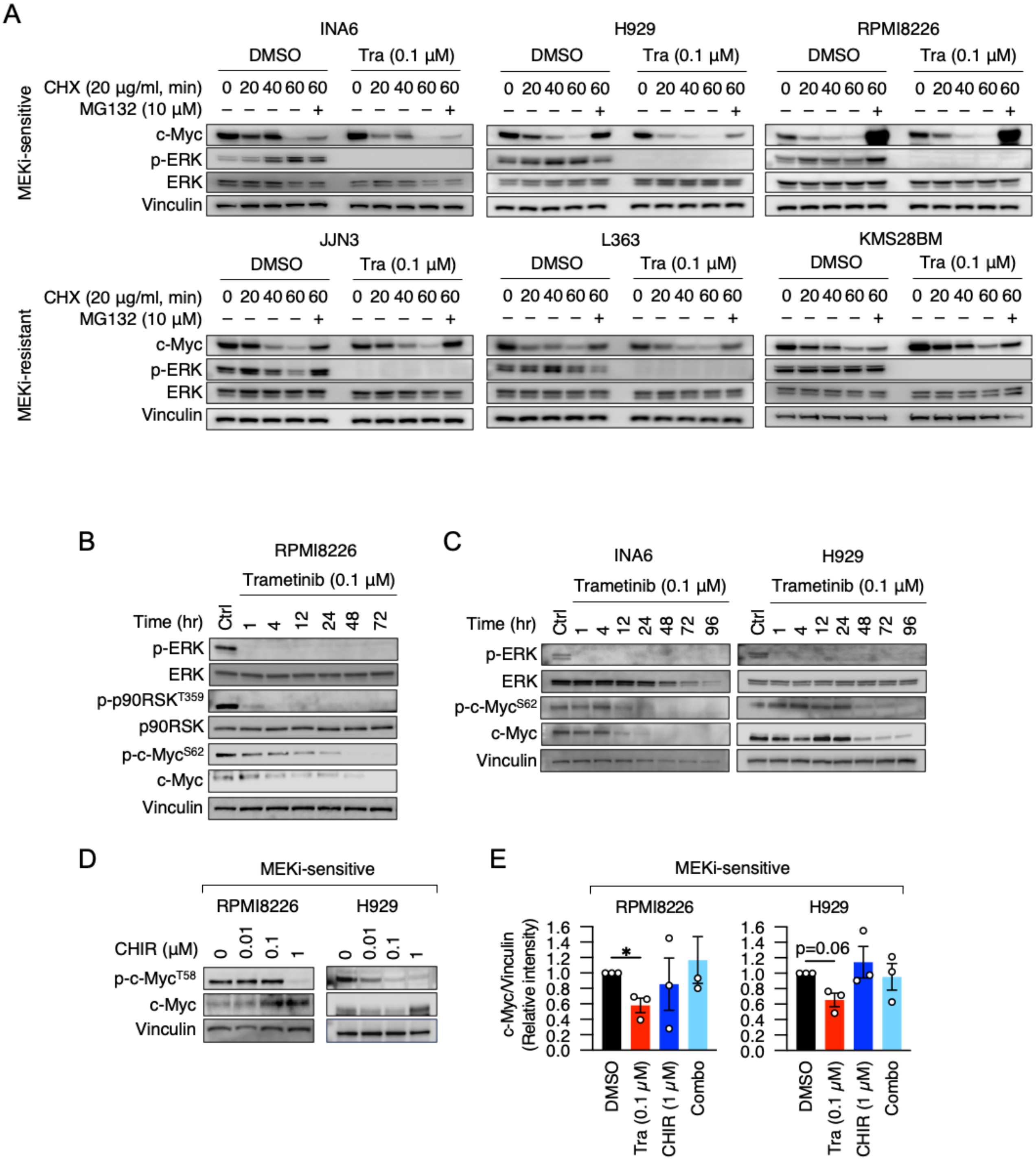
A
MEKi-sensitive
MEKi-resistant
INA6
H929
RPMI8226
JJN3
L363
KMS28BM
DMSO
Tra (0.1 µM)
CHX (20 µg/ml, min)
MG132 (10 µM)
c-Myc
p-ERK
ERK
Vinculin
B
RPMI8226
Trametinib (0.1 µM)
Time (hr)
Ctrl 1 4 12 24 48 72
p-p90RSK^T359
p90RSK
p-c-Myc^S62
C
INA6
H929
Ctrl 1 4 12 24 48 72 96
D
MEKi-sensitive
CHIR (µM)
0 0.01 0.1 1
p-c-Myc^T58
E
c-Myc/Vinculin (Relative intensity)
p=0.06
DMSO
Tra (0.1 µM)
CHIR (1 µM)
Combo

### Figure S6

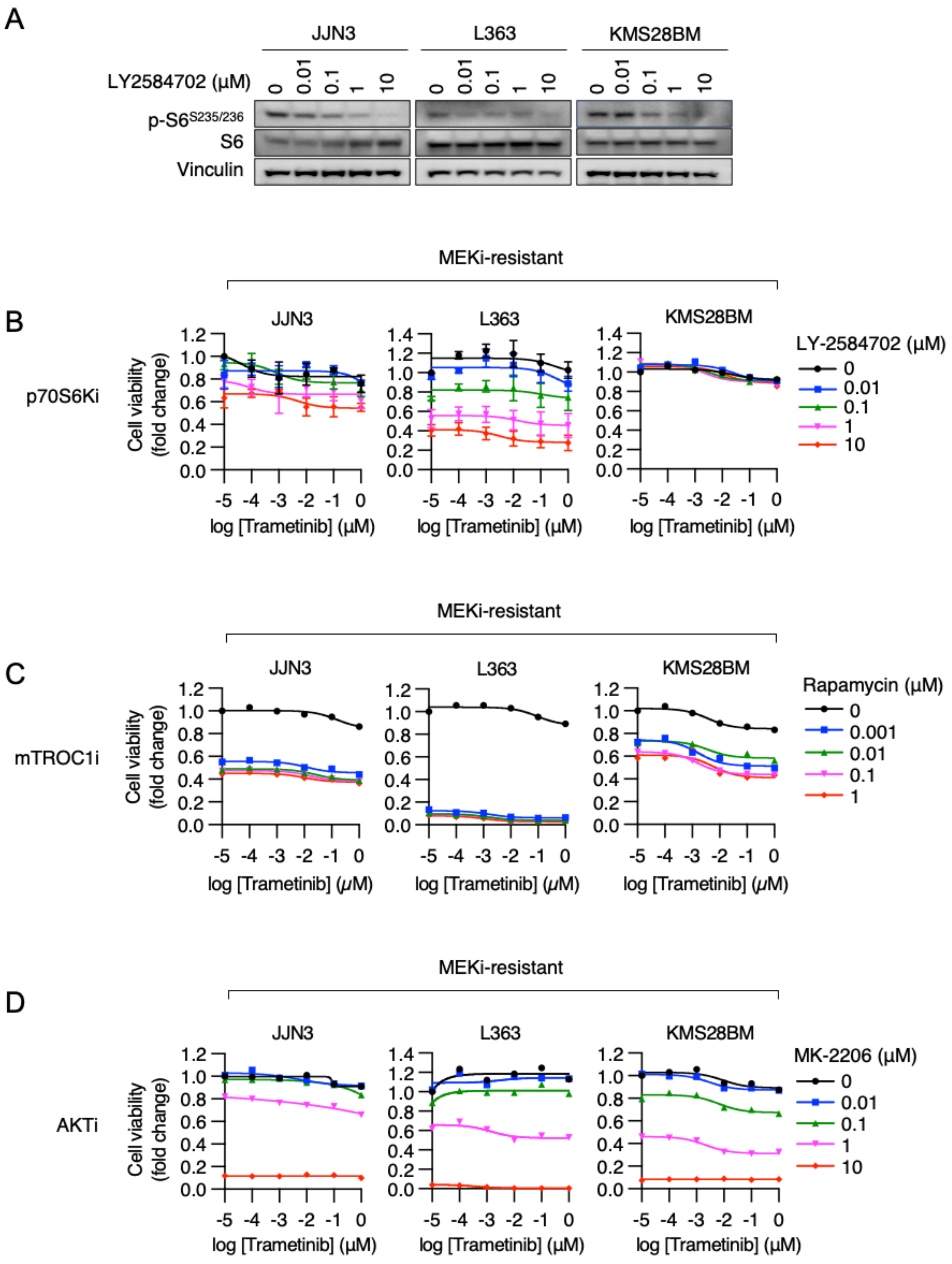
A
JJN3
L363
KMS28BM
LY2584702 (μM)
0 0.01 0.1 1 10
p-S6S235/236
S6
Vinculin
MEKi-resistant
B
p70S6Ki
Cell viability (fold change)
log [Trametinib] (μM)
LY-2584702 (μM)
0
0.01
0.1
1
10
C
mTROC1i
Rapamycin (μM)
0
0.001
0.01
0.1
1
D
AKTi
MK-2206 (μM)
0
0.01
0.1
1
10

### Figure S7

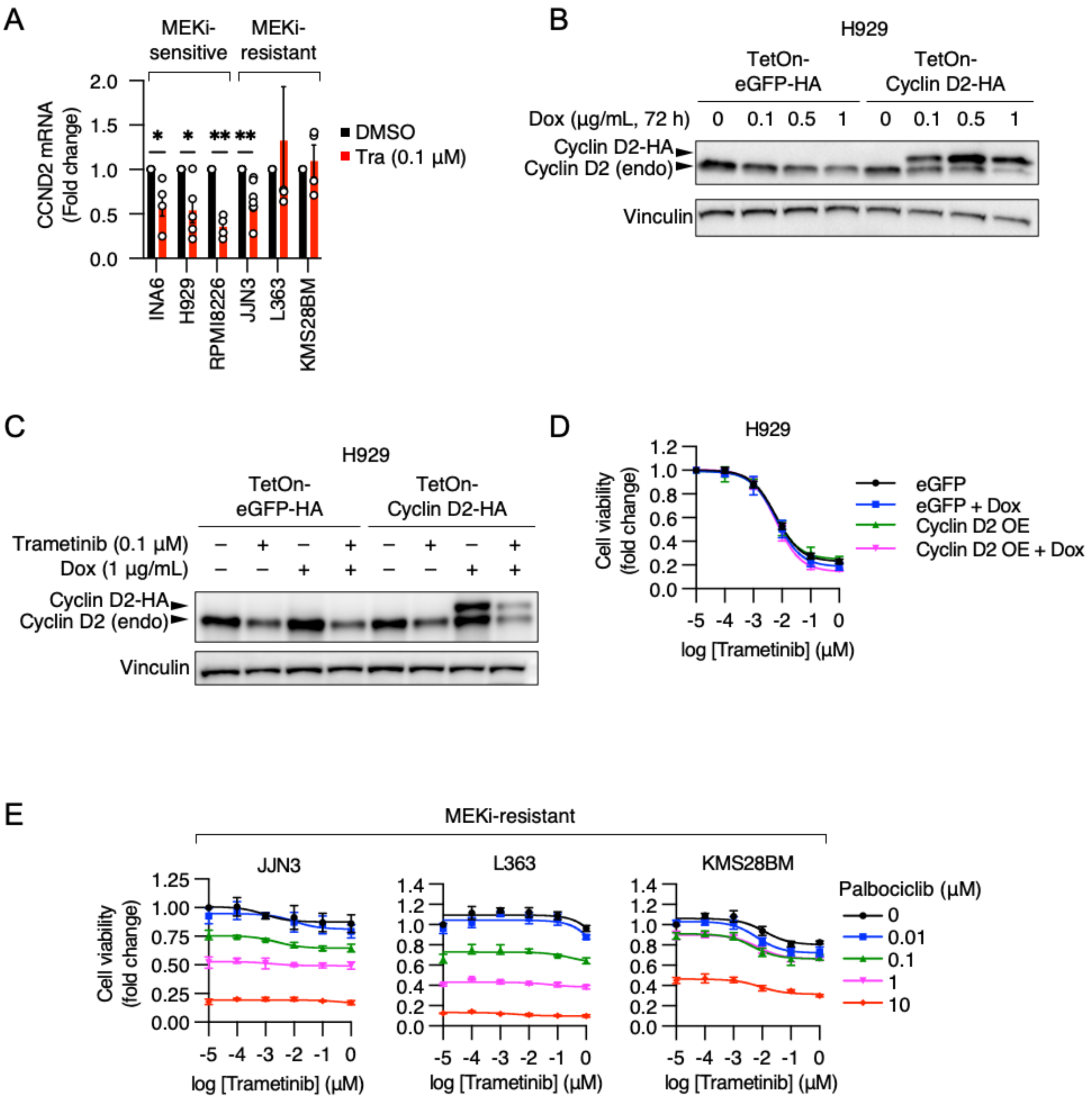
A
MEKi-sensitive
MEKi-resistant
CCND2 mRNA (Fold change)
2.0
1.5
1.0
0.5
0.0
*
*
**
**
DMSO
Tra (0.1 µM)
INA6
H929
RPMI8226
JJN3
L363
KMS28BM
B
H929
TetOn-eGFP-HA
TetOn-Cyclin D2-HA
Dox (µg/mL, 72 h)
0
0.1
0.5
1
0
0.1
0.5
1
Cyclin D2-HA
Cyclin D2 (endo)
Vinculin
C
H929
TetOn-eGFP-HA
TetOn-Cyclin D2-HA
Trametinib (0.1 µM)
Dox (1 µg/mL)
Cyclin D2-HA
Cyclin D2 (endo)
Vinculin
D
H929
Cell viability (fold change)
1.2
1.0
0.8
0.6
0.4
0.2
0.0
-5
-4
-3
-2
-1
0
log [Trametinib] (µM)
eGFP
eGFP + Dox
Cyclin D2 OE
Cyclin D2 OE + Dox
E
MEKi-resistant
JJN3
L363
KMS28BM
Cell viability (fold change)
1.25
1.00
0.75
0.50
0.25
0.00
1.4
1.2
1.0
0.8
0.6
0.4
0.2
0.0
-5
-4
-3
-2
-1
0
log [Trametinib] (µM)
Palbociclib (µM)
0
0.01
0.1
1
10

### Figure S8

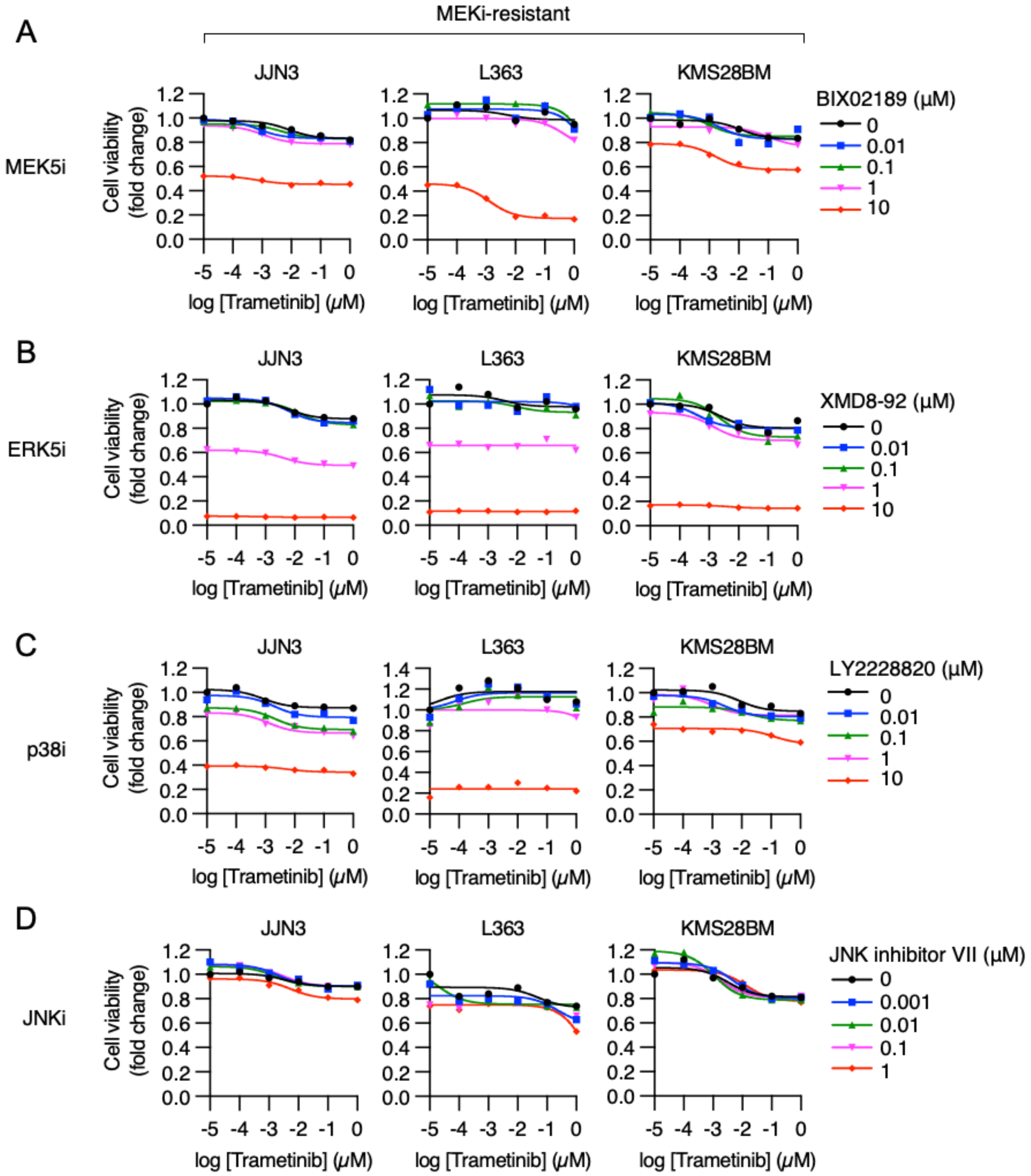
MEKi-resistant
A
B
C
D
MEK5i
ERK5i
p38i
JNKi
JJN3
L363
KMS28BM
Cell viability (fold change)
log [Trametinib] (μM)
BIX02189 (μM)
XMD8-92 (μM)
LY2228820 (μM)
JNK inhibitor VII (μM)
0
0.001
0.01
0.1
1
10

### Figure S9

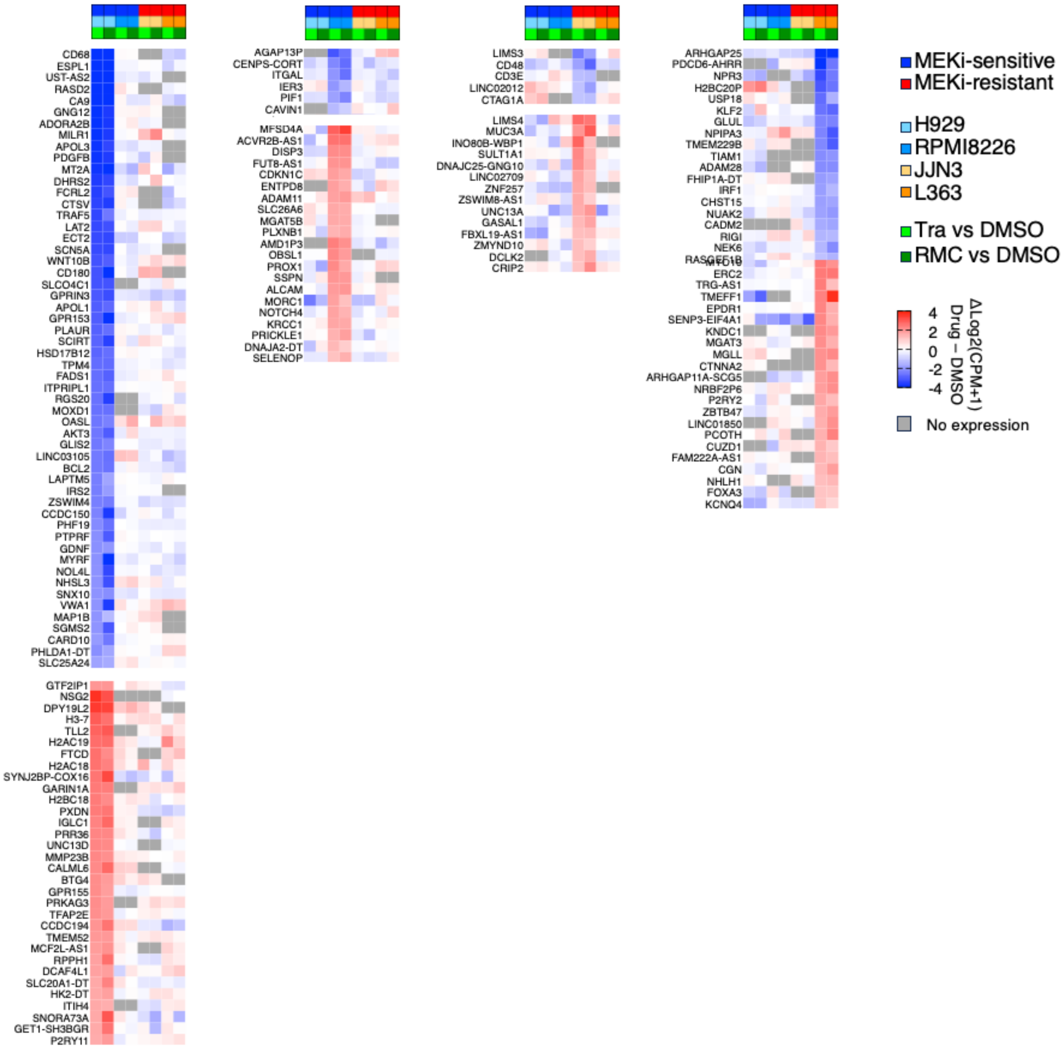
MEKi-sensitive
MEKi-resistant
H929
RPMI8226
JJN3
L363
Tra vs DMSO
RMC vs DMSO
ΔLog2(CPM+1) Drug − DMSO
4
2
0
-2
-4
No expression
CD68
ESPL1
UST-AS2
RASD2
CA9
GNG12
ADORA2B
MILR1
APOL3
PDGFB
MT2A
DHRS2
FCRL2
CTSV
TRAF5
LAT2
ECT2
SCN5A
WNT10B
CD180
SLCO4C1
GPRIN3
APOL1
GPR153
PLAUR
SCIRT
HSD17B12
TPM4
FADS1
ITPRIPL1
RGS20
MOXD1
OASL
AKT3
GLIS2
LINC03105
BCL2
LAPTM5
IRS2
ZSWIM4
CCDC150
PHF19
PTPRF
GDNF
MYRF
NOL4L
NHSL3
SNX10
VWA1
MAP1B
SGMS2
CARD10
PHLDA1-DT
SLC25A24
GTF2IP1
NSG2
DPY19L2
H3-7
TLL2
H2AC19
FTCD
H2AC18
SYNJ2BP-COX16
GARIN1A
H2BC18
PXDN
IGLC1
PRR36
UNC13D
MMP23B
CALML6
BTG4
GPR155
PRKAG3
TFAP2E
CCDC194
TMEM52
MCF2L-AS1
RPPH1
DCAF4L1
SLC20A1-DT
HK2-DT
ITIH4
SNORA73A
GET1-SH3BGR
P2RY11
AGAP13P
CENPS-CORT
ITGAL
IER3
PIF1
CAVIN1
MFSD4A
ACVR2B-AS1
DISP3
FUT8-AS1
CDKN1C
ENTPD8
ADAM11
SLC26A6
MGAT5B
PLXNB1
AMD1P3
OBSL1
PROX1
SSPN
ALCAM
MORC1
NOTCH4
KRCC1
PRICKLE1
DNAJA2-DT
SELENOP
LIMS3
CD48
CD3E
LINC02012
CTAG1A
LIMS4
MUC3A
INO80B-WBP1
SULT1A1
DNAJC25-GNG10
LINC02709
ZNF257
ZSWIM8-AS1
UNC13A
GASAL1
FBXL19-AS1
ZMYND10
DCLK2
CRIP2
ARHGAP25
PDCD6-AHRR
NPR3
H2BC20P
USP18
KLF2
GLUL
NPIPA3
TMEM229B
TIAM1
ADAM28
FHIP1A-DT
IRF1
CHST15
NUAK2
CADM2
RIGI
NEK6
RASGEF1B
ERC2
TRG-AS1
TMEFF1
EPDR1
SENP3-EIF4A1
KNDC1
MGAT3
MGLL
CTNNA2
ARHGAP11A-SCG5
NRBF2P6
P2RY2
ZBTB47
LINC01850
PCOTH
CUZD1
FAM222A-AS1
CGN
NHLH1
FOXA3
KCNQ4
